# Hunter effort increases willow ptarmigan mortality, but the cost is not visible in abundance: evaluating effort thresholds for harvest management

**DOI:** 10.64898/2026.08.27.747286

**Authors:** Tomas Willebrand, Maria Hörnell-Willebrand, Rolf Brittas, Eivind Kleiven

## Abstract

Managers must often regulate harvest under uncertainty, and thresholds that trigger action when exceeded offer a practical way to manage priorities. In 2005, guidelines for willow ptarmigan (*Lagopus lagopus*) harvest on state-managed land in Sweden recommended that hunter effort be kept around one hunter-day km^−2^ and not exceed three hunter-days km^−2^. We evaluated these recommendations using line-transect counts and harvest data from six sites, forming three pairs of hunted and closed sites, over twelve years (1996–2007). We fitted a distance-sampling model of adult and juvenile density and two dynamic models: a Hazard model linking effort to harvest mortality through the reported bag, and a Gompertz model of population growth.

Densities fluctuated strongly between years, largely in synchrony across all six sites regardless of hunting. In the Hazard model, harvest mortality increased clearly with effort. The per-bird harvest probability was 6% at one hunter-day km^−2^, 18% at three and about 36% at the highest observed effort (6.6 hunter-days km^−2^), reducing annual survival from 0.40 without harvest to 0.37 and 0.32 at one and three hunter-days km^−2^, respectively. In the Gompertz model, annual population change was driven mainly by breeding success around site-specific levels, and the effect of effort on growth was not resolved. Hunted sites held more birds than predicted from survival. This local compensation, through immigration or lower natural mortality, offset part of the harvest loss but left net reductions of about 4% at low and 9% at high effort. Harvest therefore reduces survival, but at the observed effort levels its consequences were not detectable in abundance data alone.

A fixed effort limit, anchored to monitored effort and bag, is an adequate design given the information available to managers, because it rests on the relationship this study establishes firmly: the translation of effort into harvest mortality. Our results support the existing recommendations and argue for annual follow-up of effort alongside the population counts.

## 1. Introduction

Managers of harvested populations must act under uncertainty, often with scarce and noisy data and incomplete understanding of what drives population change (Regan et al. 2002; Walsh et al. 2015). Adaptive management, in which the population model is updated as data accumulate, is the established solution (Walters, 1986; Nichols et al. 2007), but resources rarely allow it for most harvested species.

A simpler alternative is threshold management, in which action is taken when a limit is exceeded (Lande et al. 1997). The thresholds evaluated here are of the simplest kind: fixed limits on hunter effort, which require no annual population estimate.

In the early 1990s in Sweden, most of the state-managed tundra, about 60 000 km^2^, was made accessible to sport hunting (Sveriges riksdag, 1993). The reform prompted discussions on how to best track the willow ptarmigan (*Lagopus lagopus*) population, and the use of effort and bag limits to secure a sustainable harvest (Eriksson et al. 2006). By 1996, about twenty sites had been established to monitor regional willow ptarmigan populations, providing estimates of adult numbers and reproductive success. In 2005, early analyses of a crude relationship between effort, harvest rate and population response recommended that harvest effort should be kept around one hunter-day km^−2^ and should not exceed three hunter-days km^−2^ (Eriksson et al. 2006).

Reproductive success and recruitment show large annual fluctuations (Steen, Steen, et al. 1988; Hörnell-Willebrand, Marcström, et al. 2006). These seemingly stochastic processes, and the presence of complex time-lags (Tornberg, Reif, et al. 2012; Turchin, 2003), have made the development of management models problematic. Predation is the primary cause of nest loss and of chick and adult mortality, and *per capita* reproductive success in autumn tends to be positively correlated with the abundance of microtine rodents (Steen and Haugvold, 2009). In addition, variation in the onset of spring and in plant production affects the nutritional condition of breeding females and subsequent breeding success (Brittas, 1988; Willebrand, Hörnell-Willebrand, et al. 2025; Eriksen et al. 2025). Analyses of willow ptarmigan populations have found moderate but significant negative density dependence (Pedersen et al. 2004; Willebrand, Hörnell-Willebrand, et al. 2025), and compensatory predation and immigration through natal dispersal are the most likely mechanisms. Because *per capita* reproductive success does not depend on adult density (Hörnell-Willebrand, Marcström, et al. 2006), density dependence must act through survival or dispersal. A reduced level of natural mortality as a compensatory response to harvest mortality has been addressed in several studies, but the conclusions about the strength of compensatory mortality remain ambiguous (Smith and Willebrand, 1999; Willebrand and Hörnell, 2001; Pedersen et al. 2004; Israelsen et al. 2020; Sandercock et al. 2011). Using radio-collared willow ptarmigan, Smith and Willebrand (1999) estimated harvest mortality to be almost completely additive compared with a closed site, but did not detect the expected difference in population development between the open and closed sites. Later work proposed that long-distance dispersal of juvenile females reduced the difference in breeding density (Hörnell-Willebrand, Willebrand, et al. 2014).

If harvest effects can be detected in survival but are masked in abundance, evaluating effort thresholds requires models that address both. Here we evaluate 12 years of population development at six sites forming three pairs of sites open and closed to sport harvest. Our aims were to 1) re-evaluate the previous threshold recommendations of 1 and 3 accumulated hunter-days km^−2^, 2) estimate the effects of harvest on population development, and 3) make management recommendations. We combined hunter effort and bag records with distance-sampling data from 1996–2007 in a static model of population density, which fed two dynamic models: a mechanistic Hazard model that links effort to harvest mortality, and a phenomenological Gompertz model of population growth.

## 2. Methods

### 2.1. Study area

In the southern part of the tundra region of Sweden, the state-managed area of Jämtland County is about 10 600 km^2^, divided into 113 management units with an average size of 94 km^2^. The overall landscape consists of three-quarters alpine heath and shrubland, and one-quarter mountain birch with scattered patches of conifers. Common predators of willow ptarmigan are gyrfalcon *Falco rusticolus*, goshawk *Accipiter gentilis*, red fox *Vulpes vulpes*, and stoat *Mustela erminea*. The red fox and stoat show both functional and lagged numerical responses to fluctuating small-rodent populations (Rød-Eriksen et al. 2023). In this study, we used data from six sites forming three open/closed pairs (Figure 1). The sites were chosen to represent the overall habitat types of the managed area in 1994/95. The total area was 279.3 km^2^ for open sites (A1 = 71.7, B1 = 164.0, C1 = 43.6 km^2^) and 326.4 km^2^ for closed sites (A0 = 73.9, B0 = 177.7, C0 = 74.8 km^2^). The open sites were subject to the same hunting regulations as all open sites in the county.

**Figure 1:**
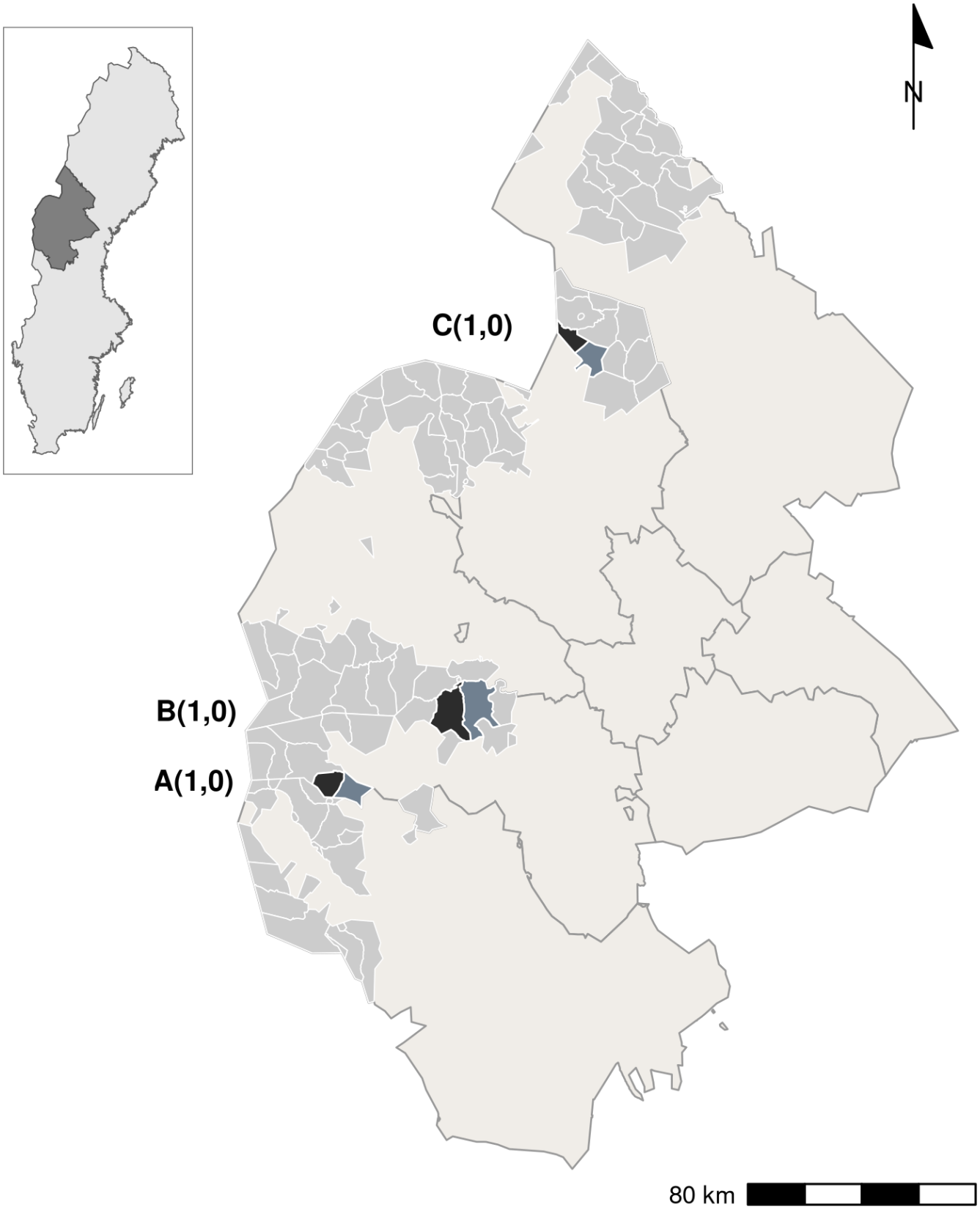
Study sites in the state-managed mountain area of Jämtland County, central Sweden. The six sites form three pairs (A, B, C), each with an open (hunted; 1, black) and a closed (unhunted; 0, blue-grey) site. Light grey areas are the other state-managed small-game management units, all open to hunting; the remainder of the county is shown in beige. Inset: location of Jämtland County (dark grey) in Sweden.

### 2.2. Field data

#### Line transects

Between 1996 and 2007, adult and young willow ptarmigan were counted in early August, 2–4 weeks before the hunting season. Transect lines covered all sites below 1100 m a.s.l.; lines were placed 400 m apart and oriented from low to high elevations. Total transect length per site and year varied from 56 to 101 km (mean 84 km; Table 4 in Appendix A). Encounter data were prepared from the original field records: transect lines without observations were kept as survey effort, birds of unknown age (6 % of all birds) were assigned to adults and young from the composition of their group, and groups of young seen without an adult were treated as broods with one unseen adult. Pointing dogs were used to detect willow ptarmigan at different distances from the line. The dog handlers were selectively recruited and received annual feedback. A census team of dog handlers counted both sites in a pair on the same occasion. Group size, number of adults and young, and perpendicular distance to the line were recorded for each encounter. We registered 3 750 group encounters, with between 147 and 420 encounters per year.

#### Bag records

The hunting season ran from 25 August to the end of February. Willow ptarmigan were hunted predominantly with shotguns over pointing dogs, and approximately two-thirds of all effort occurred during the first ten days (Eriksson et al. 2006), before brood breakup and dispersal (Smith and Willebrand, 1999). All licensed hunters could obtain a hunting permit; foreign hunters were not allowed to hunt from 1996 to 1998. Effort and bag records (days hunted and birds killed) were reported within one week of each trip, with 70–96 % compliance. Records were adjusted for non-response, assuming non-respondents did not differ systematically from respondents. Effort was defined as total hunter-days (hd) and bag size as the total number of willow ptarmigan shot. Annual harvest data for each management site are recorded by the regional county administrations, and we used a preliminary analysis of the regional hunter effort in all sites for 2024/2025 to estimate any change from our study period 1996–2007.

### 2.3. Statistical models

#### Rationale and overall approach

We developed three sets of Bayesian hierarchical models fitted by MCMC using JAGS (Plummer, 2003). The static model provides the densities of adults and young; its detection estimates are passed to two dynamic models that re-estimate density as a latent state. The Hazard model splits annual survival into natural and harvest hazards and links harvest to effort and the reported bag; it answers at what effort hunting increases mortality. The Gompertz model describes population growth as a function of density, productivity and effort; it asks whether an effort effect is visible in abundance. Model schematic in Figure 4 (Appendix A).

#### Static distance-sampling model

Detection probability at perpendicular distance *x* from the transect was described by the hazard-rate function:

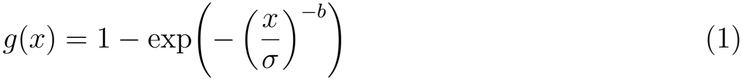

where *σ* is the scale parameter and *b* the shape parameter. The shape parameter was fixed separately for brood and non-brood encounters based on values from preliminary model runs; the scale *σ* varied across sites, years, and encounter types through a hierarchical structure with site and year random effects. Encounter counts at each site and year followed a Poisson distribution with expected value proportional to group density, effective surveyed area, and detection probability. Adult group size (1, 2, or 3+ birds) was modeled via a hierarchical multinomial logit model. The number of young per group was modeled with a two-part structure: a logistic model for brood occurrence and a zero-truncated negative binomial for positive brood sizes. Full prior specifications and model details are in Appendix A.

Detection uncertainty was propagated into the dynamic models without refitting the full observation model. The posterior of logit-scale detection probability at each site-year was summarized as a normal distribution (mean and standard deviation) and used as an informative prior for detection probability in the dynamic models.

#### Hazard-based survival dynamics

Annual (August-to-August) survival at site *s* in year *t* is the product of two competing constant-hazard processes:

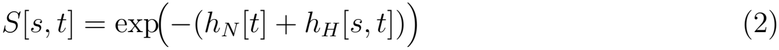

where *h*_*N*_ [*t*] is the natural mortality hazard shared across sites in year *t*, and *h*_*H*_[*s, t*] is the harvest hazard (zero for closed sites). In the standardized model, effort enters as *E*_*sc*_ = (log(1 + intensity) − *µ*_*I*_)*/σ*_*I*_ on a per-10 km^2^ intensity scale; the standardization center (*E*_*sc*_ = 0) corresponds to 1.24 hd km^−2^, and the geometric mean (1.20 hd km^−2^) corresponds to *E*_*sc*_ ≈ −0.04. The harvest hazard is linked to standardized hunter effort through a log-linear model with a year-level random effect that separates year-to-year variation in hunting efficiency from the effort effect of primary interest:

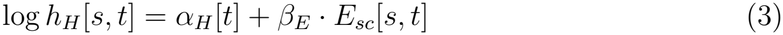

Next year’s adult density is modeled as:

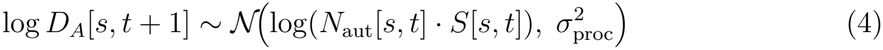

where *N*_aut_[*s, t*] = *D*_*A*_[*s, t*] + *D*_*Y*_ [*s, t*] is total autumn density and 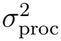 captures process noise including demographic stochasticity and net dispersal. *The annual bag is linked to the hazard parameters through a negative binomial likelihood, so that bag counts provide an independent constraint on the harvest hazard*, with juveniles weighted by an availability ratio *κ*_*Y*_ ; further details are given in Appendix A.

The natural hazard *h*_*N*_ [*t*] is shared by open and closed sites, so closed sites anchor natural mortality. With the additive form of Eq. 2 the model cannot represent compensation, which is absorbed by the process error; we therefore interpret *h*_*H*_ as gross harvest mortality and assess compensation separately (paragraph 2.3). Because *h*_*N*_ is estimated from apparent survival, it includes net dispersal.

#### Gompertz population dynamics

Log adult density follows an autoregressive process:

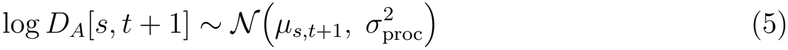

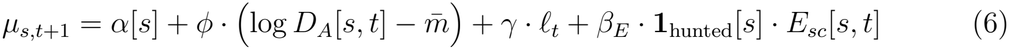

where *α*[*s*] is the site intercept (the expected log density in year *t* + 1 when current density equals 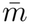), *ϕ* ∈ (0, 1) is the autoregressive coefficient constrained by a logistic-transformed prior (*ϕ* = 1 means no density dependence, and 1 − *ϕ* is the proportion of a deviation from the site’s equilibrium level removed per year), 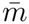 is the grand mean log-density used to center the density covariate, *ℓ*_*t*_ is the productivity index 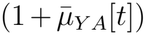) in year *t*, centered on its mean over years, and *β*_*E*_ is the harvest effort coefficient. The site intercept is built hierarchically with pair-level and site-level components to respect the paired design (see Appendix A4).

#### Model validation

Priors were weakly informative, and posteriors were checked for accumulation at prior boundaries. Final runs used 12 chains of 18 000 iterations after 4 000 burn-in (168 000 draws); convergence was assessed by 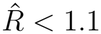 and effective sample size *>* 200. Model fit was evaluated with posterior predictive checks; results are in Appendix A6.

#### Harvest, survival cost, and population consequences

We explored the relationship between effort and harvest rate, defined as the per-bird harvest probability *p*_*H*_ = 1 − exp(−*h*_*H*_), using the full posterior of the mean harvest-hazard intercept *α*_*H,µ*_ and the effort slope *β*_*E*_. Absolute annual survival was evaluated at the posterior mean natural hazard 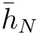 (the mean over years of *h*_*N*_ [*t*]) plus the effort-specific harvest hazard. The harvest-only survival factor *S*_*h*_, the harvest probability *p*_*H*_ and the harvest-attributable reduction in annual survival Δ*S* were computed per posterior draw as functions of standardized hunter effort:

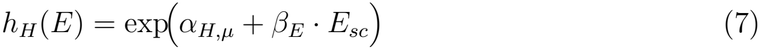

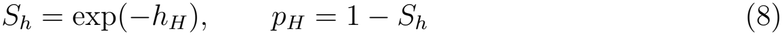

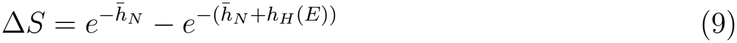

#### What the Gompertz model can and cannot resolve regarding effort

The Gompertz effort coefficient is identified only from year-to-year variation in effort within the three open sites, and *ϕ* is weakly identified from eleven transitions per site (Appendix A4). We therefore report the Gompertz model as a description of short-term dynamics, not as a basis for effort thresholds.

#### Local compensation of harvest mortality

We use compensation for any process that makes the population effect of harvest smaller than the harvest loss. Four mechanisms are possible: lower natural mortality when harvest reduces density, net immigration to hunted sites, a delayed reduction in natural loss in the following year, and buffering by unhunted surroundings at the regional scale. The Hazard model excludes the first by construction, and the Gompertz model cannot estimate it, because its density feedback is largely determined by the prior. The residual contrast below measures the first two together at the scale of a site pair; we call this local compensation. The last two are not addressed.

In the Hazard model, harvest mortality is additive by construction (Eq. 2), and the natural hazard *h*_*N*_ [*t*] is shared by open and closed sites. We examined whether hunted sites had more birds than predicted by the additive model, and used the posterior draws of the already fitted Hazard model to evaluate the exchange of birds between a hunted site and its surrounding area, as proposed by Hörnell-Willebrand, Willebrand, et al. (2014). This approach works at the local scale, but whether harvest is compensated at large regional scales cannot be evaluated with six sites. We return to compensation at the regional scale in the Discussion.

For each posterior draw we computed the residual of the process equation,

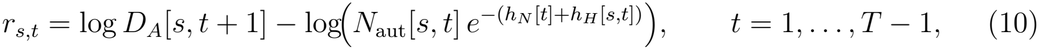

the log ratio of the adult density present in year *t* + 1 to the density expected under additive mortality. Because *h*_*N*_ [*t*] is estimated jointly from all six sites, a systematic excess on open sites is partly absorbed by a lower *h*_*N*_ [*t*] and then appears as a deficit on closed sites. Instead of interpreting the residuals directly, we contrasted each open-site residual with the mean residual of the three closed sites in the same year,

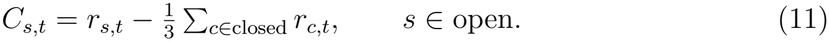

The within-year contrast removes unusual conditions shared by all sites in a year, and prevents the clustering of high effort in 2003–2007 from being confounded with year. Each draw gives 33 contrasts (three open sites × 11 transitions). A positive *C*_*s,t*_ means that a hunted site gained or retained more birds than the closed sites, relative to the additive prediction.

We summarized the posterior of the mean contrast 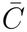 over all open site-years, per pair, and separately for site-years above and below the standardization effort center (*E*_*sc*_ = 0, 1.24 hd km^−2^), and reported it as a percentage, 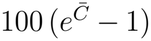. Within each effort class we combined the gain with the harvest loss. With *L* the mean harvest probability 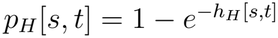 over the site-years in the class, the net effect of harvest relative to the closed sites is

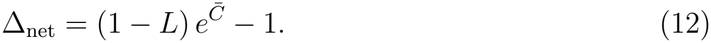

Negative values mean that the harvest loss is not fully offset. We report the posterior probability of full compensation, *P* (Δ_net_ ≥ 0).

## 3. Results

### 3.1. Distance sampling and densities

The distance-sampling model separated adult and young densities. Adult and juvenile densities showed pronounced year-to-year variation that was largely synchronous across all six sites regardless of hunting status (Figure 2). Juvenile densities were consistently more variable than adult densities and showed the most extreme fluctuations in years of poor and good productivity. Across all six sites during the 12-year study period (1996–2007), adult density ranged from approximately 3 to 11 ind. km^−2^, and juvenile density showed greater absolute variation, reaching up to 25 ind. km^−2^. Annual productivity (young per adult, 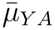) ranged from 1.40 (1996) to 2.26 (2003). Detection probability varied among site-years (0.30–0.62; Appendix A2).

**Figure 2:**
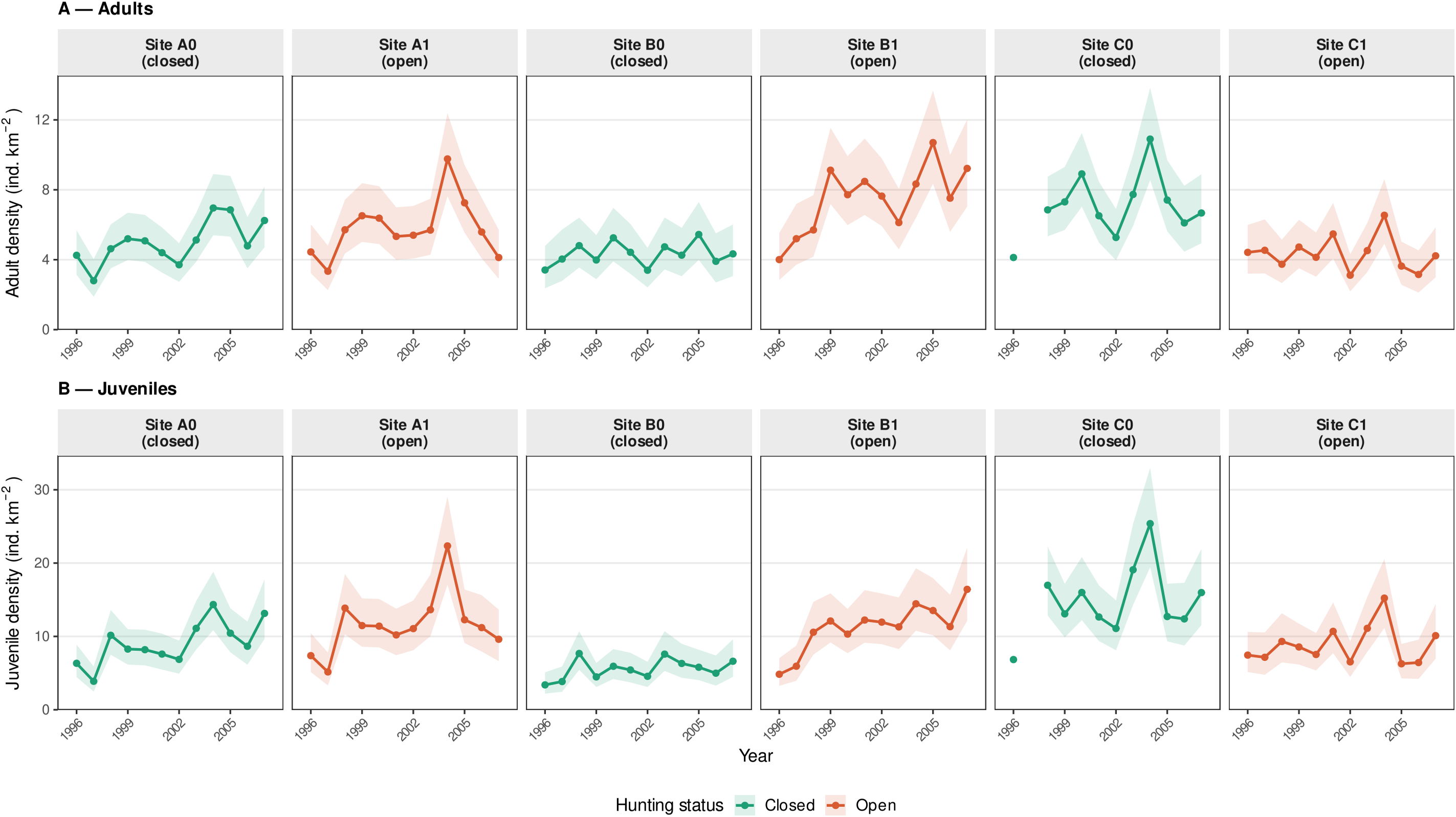
Density (ind. km^−2^) of (A) adults and (B) juveniles on the six study sites in August, before the hunting season, 1996–2007, estimated by the static hierarchical distance-sampling model. Lines show posterior medians and shaded ribbons 95 % credible intervals. Letters denote site pairs; 0 = closed (unhunted, teal) and 1 = open (hunted, orange). Adult and juvenile densities are derived jointly from the estimated density of groups and the modeled number of adults and young per group. Site C0 was not surveyed in 1997.

### 3.2. Posterior parameter estimates of dynamic models

Here we present posterior summaries of key biological parameters from both models. All parameters converged with Gelman–Rubin statistics 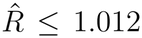 and are presented in Table 1. The harvest effort parameter *β*_*E*_ is estimated on different scales in the two models and with very different precision. In the Hazard model, *β*_*E*_ = 0.798 (95 % CrI 0.627–0.954, *f* = 1.000) was positive and excluded zero with near certainty, supporting the hypothesis that higher hunter effort increases mortality hazard. An increase in effort of one standard deviation above the center of the standardized scale (from 1.24 to 2.73 hd km^−2^) reduced annual survival on hunted sites by 4.0 percentage points (Δ*S* = −0.040, 95 % CrI −0.067 to −0.021), and harvest survival at +1 SD effort was 0.891 (95 % CrI 0.812–0.945) of that at the center of the scale. In the Gompertz model, the parameter was *β*_*E*_ = −0.020 (95 % CrI −0.095 to +0.058, *f* = 0.696), and pointed in the same direction as the Hazard model (higher effort, lower growth), but its credible interval included zero.

**Table 1.**
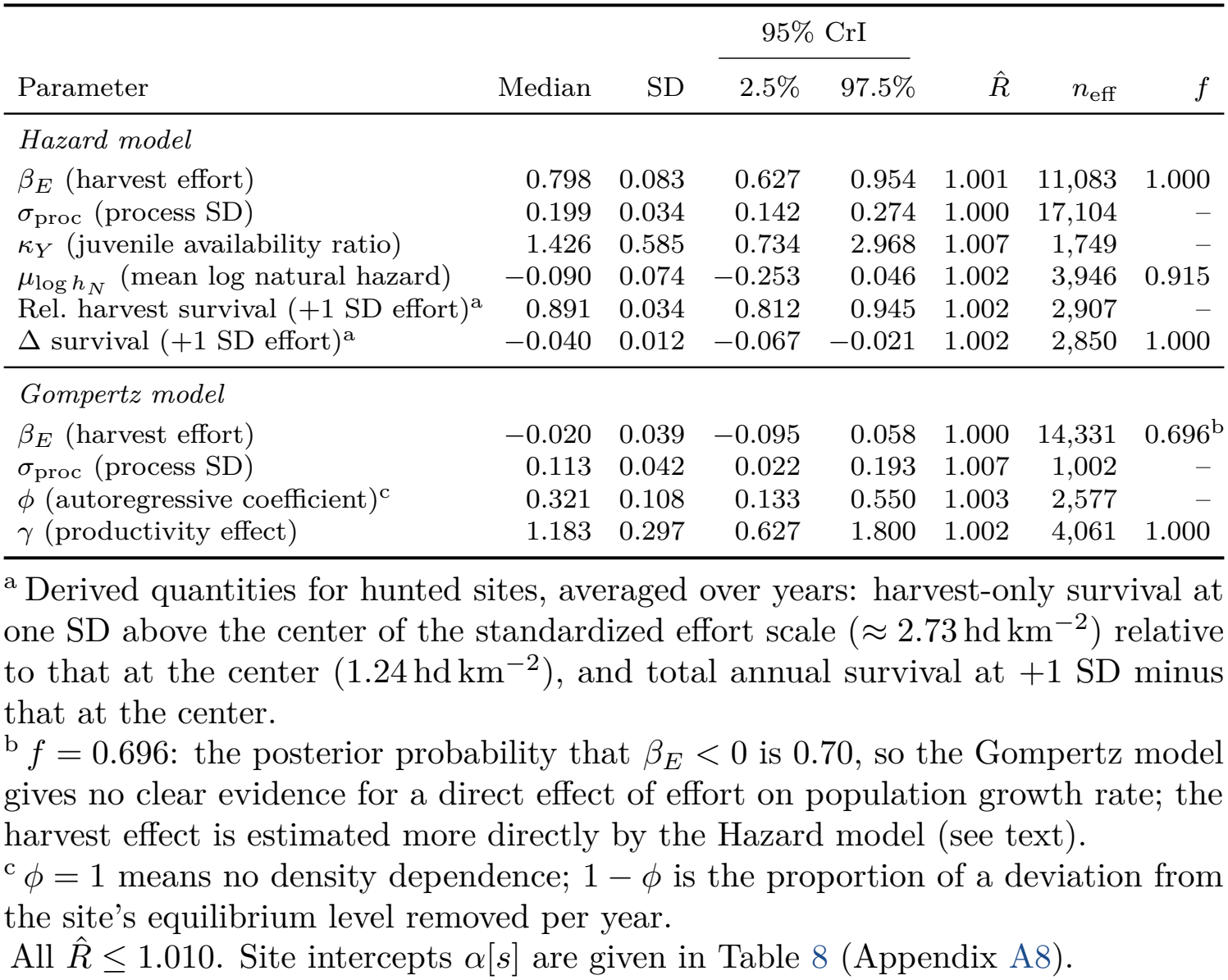
Posterior summaries of key parameters from the Hazard and Gompertz state-space models: posterior median, standard deviation (SD), 95 % credible interval (CrI), Gelman–Rubin diagnostic 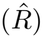, effective sample size (*n*_eff_), and *f*, the proportion of the posterior with the same sign as the posterior mean. *f* is not shown for quantities that are positive by definition. Based on 168,000 posterior draws from 12 chains; full summaries in Tables 7 and 8.

|  |  |  | 95% CrI |  |  |  |  |
|---|---|---|---|---|---|---|---|
| Parameter | Median | SD | 2.5% | 97.5% | R̂ | n<sub>eff</sub> | *f* |
| *Hazard model* |  |  |  |  |  |  |  |
| β<sub>E</sub> (harvest effort) | 0.798 | 0.083 | 0.627 | 0.954 | 1.001 | 11,083 | 1.000 |
| σ<sub>proc</sub> (process SD) | 0.199 | 0.034 | 0.142 | 0.274 | 1.000 | 17,104 | – |
| κ<sub>Y</sub> (juvenile availability ratio) | 1.426 | 0.585 | 0.734 | 2.968 | 1.007 | 1,749 | – |
| μ<sub>log h<sub>N</sub></sub> (mean log natural hazard) | −0.090 | 0.074 | −0.253 | 0.046 | 1.002 | 3,946 | 0.915 |
| Rel. harvest survival (+1 SD effort)<sup>a</sup> | 0.891 | 0.034 | 0.812 | 0.945 | 1.002 | 2,907 | – |
| Δ survival (+1 SD effort)<sup>a</sup> | −0.040 | 0.012 | −0.067 | −0.021 | 1.002 | 2,850 | 1.000 |
| *Gompertz model* |  |  |  |  |  |  |  |
| β<sub>E</sub> (harvest effort) | −0.020 | 0.039 | −0.095 | 0.058 | 1.000 | 14,331 | 0.696<sup>b</sup> |
| σ<sub>proc</sub> (process SD) | 0.113 | 0.042 | 0.022 | 0.193 | 1.007 | 1,002 | – |
| ϕ (autoregressive coefficient)<sup>c</sup> | 0.321 | 0.108 | 0.133 | 0.550 | 1.003 | 2,577 | – |
| γ (productivity effect) | 1.183 | 0.297 | 0.627 | 1.800 | 1.002 | 4,061 | 1.000 |

<sup>a</sup> Derived quantities for hunted sites, averaged over years: harvest-only survival at one SD above the center of the standardized effort scale (≈ 2.73 hd km<sup>−2</sup>) relative to that at the center (1.24 hd km<sup>−2</sup>), and total annual survival at +1 SD minus that at the center.
<sup>b</sup> *f* = 0.696: the posterior probability that β<sub>E</sub> < 0 is 0.70, so the Gompertz model gives no clear evidence for a direct effect of effort on population growth rate; the harvest effect is estimated more directly by the Hazard model (see text).
<sup>c</sup> ϕ = 1 means no density dependence; 1 − ϕ is the proportion of a deviation from the site's equilibrium level removed per year.
All R̂ ≤ 1.010. Site intercepts α[*s*] are given in Table 8 (Appendix A8).

In the Gompertz model, annual population change was driven mainly by productivity (*γ* = 1.183, 95 % CrI 0.627–1.800). The density-dependence parameter (*ϕ* = 0.321, 95 % CrI 0.133–0.550) was largely determined by its prior (Appendix A4). Equilibrium densities differed between open and closed sites in both directions (open relative to closed: A +18 %, B +87 %, C −44 %), so persistent differences in level cannot be attributed to harvest. The process standard deviations of the two models are compared in Appendix A6.

The natural mortality hazard (Table 1) indicates substantial baseline mortality, corresponding to annual survival of 0.40 in the absence of harvest. Mean annual survival on hunted sites, integrating both natural and harvest mortality hazards, had a posterior median of 0.365 (95 % CrI 0.342–0.390); on average, about 63 % of the autumn population on open sites did not survive or emigrated before the following breeding season.

### 3.3. Harvest effort, survival cost, and population consequences

Because the Gompertz model did not resolve an effort effect (Section 3.2), the survival costs and population consequences below are derived from the Hazard model.

Hunter effort on open sites had a geometric mean of 1.20 hd km^−2^ (range 0.16–6.57), increased over the study period with the highest values in 2004–2005, and was not significantly related to annual reproductive success (*r* = 0.19, *p* = 0.26).

The Hazard model translated the effort levels into harvest probabilities (Figure 3). At the lower management reference threshold of 1 hd km^−2^, the predicted harvest probability was 6.5 % (95 % CrI 3.6–10.0 %), meaning that about 1 in 15 birds was shot. At the upper threshold of 3 hd km^−2^, harvest probability rose to 18.2 % (95 % CrI 10.0–28.5 %), and at the upper bound of the observed effort range (6.6 hd km^−2^), it reached approximately 36 %.

**Figure 3:**
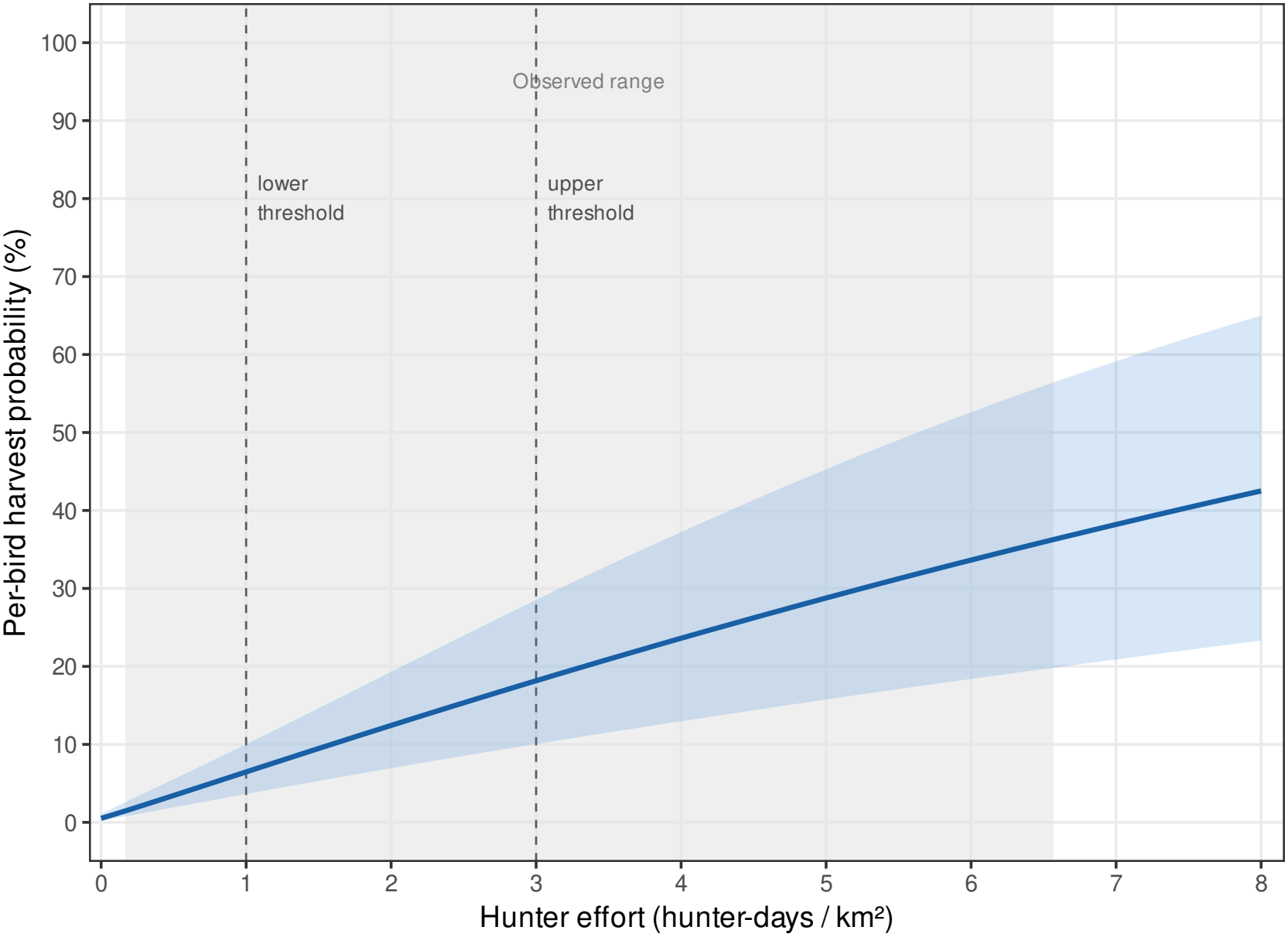
Per-bird harvest probability as a function of hunter effort (hunter-days, hd, per km^2^) for an open (hunted) site at the mean year effect (*α*_*H,µ*_), from the Hazard model. Harvest probability is the probability that a bird present in autumn is shot during the hunting season, 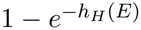. The solid line is the posterior median and the blue band the 95 % credible interval; the grey area marks the range of effort observed on open sites (0.16–6.6 hd km^−2^); dashed lines mark the lower and upper management reference thresholds (1 and 3 hd km^−2^). Predictions beyond the observed range are extrapolations.

**Figure 4:**
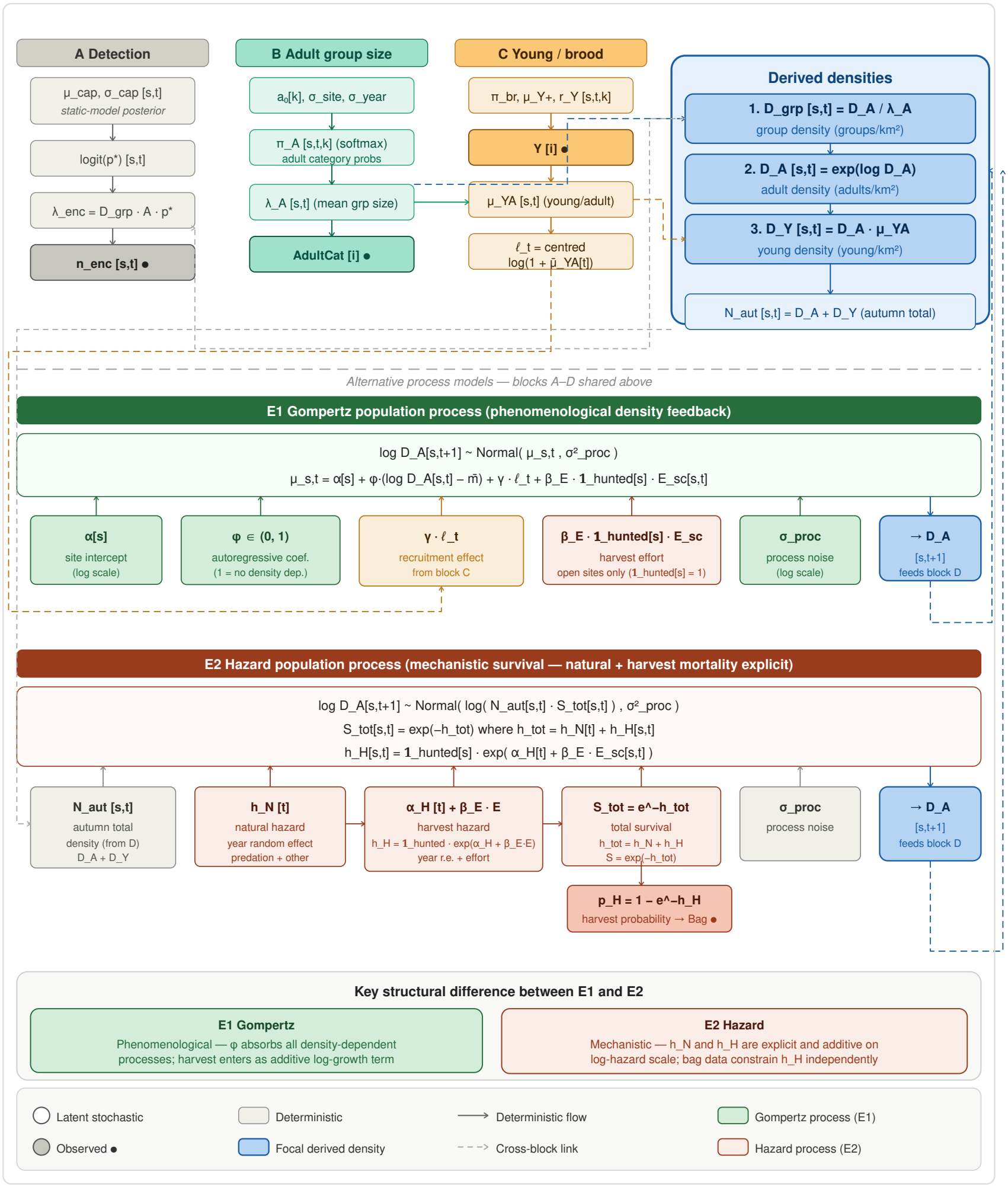
Schematic of the dynamic state-space models: the shared observation structure (A, detection; B, adult group size; C, young per brood; D, derived densities) and the two complementary process models, E1 Gompertz (green) and E2 Hazard (red). Detection in block A uses the posterior of the static distance-sampling model as an informative prior. Observed data are marked with •; beige boxes are deterministic quantities; the blue panel holds the derived densities that link the observation and process models. Solid arrows show deterministic flow, dashed arrows show links between blocks, and boxes marked → *D*_*A*_[*s, t*+1] the transition to the next year. The bottom panel summarizes the structural difference between E1 and E2.

These harvest probabilities translate into a survival cost (Table 2). Annual survival was 0.40 without harvest; harvest reduced it by 2.6 percentage points to 0.37 at 1 hd km^−2^, by 7.2 pp to 0.32 at 3 hd km^−2^, and by 14.4 pp to 0.25 at the peak effort recorded at site A1 (6.6 hd km^−2^), a 36 % proportional reduction equal to the per-bird harvest probability (Figure 3). Because the harvest and natural mortality hazards are additive, the survival cost of harvest depends only on effort, rising at a slightly decreasing rate per hunter-day, and not on productivity. Productivity instead determines how far the losses are replaced by recruitment in the following breeding season.

**Table 2.** Survival cost of harvest for willow ptarmigan on an open (hunted) site at the mean year effect (*α*_*H,µ*_) as a function of hunter effort (hunter-days, hd, per km^2^), derived from the Hazard model. The reduction is the harvest-attributable loss in annual survival relative to zero effort, in percentage points (pp); absolute survival is annual survival including harvest. Values are posterior medians with 95 % credible intervals, computed separately for each quantity per posterior draw, so the interval for the reduction cannot be obtained by subtracting the survival intervals.

| Effort (hd km<sup>−2</sup>) | Harvest-attributable survival reduction (pp), median [95 % CrI] | Absolute annual survival, median [95 % CrI] |
|---|---|---|
| 0 (no harvest) | – | 0.40 [0.37, 0.42] |
| 1.00 (lower management threshold) | 2.6 [1.4, 4.1] | 0.37 [0.35, 0.39] |
| 1.24 (center of standardized scale) | 3.1 [1.7, 5.0] | 0.36 [0.34, 0.39] |
| 2.00 | 4.9 [2.7, 7.9] | 0.35 [0.32, 0.37] |
| 3.00 (upper management threshold) | 7.2 [3.9, 11.7] | 0.32 [0.29, 0.36] |
| 5.00 | 11.4 [6.1, 18.5] | 0.28 [0.22, 0.33] |
| 6.60 (site A1 peak) | 14.4 [7.7, 23.1] | 0.25 [0.18, 0.31] |

The reduction is $\Delta S = \exp(-\bar{h}_N) - \exp(-(\bar{h}_N + h_H(E)))$, where $\bar{h}_N$ is the natural hazard averaged over years and $h_H(E)$ the harvest hazard at effort *E* for the mean year effect ($\alpha_{H,\mu}$), both evaluated per posterior draw. Survival is apparent survival, net of emigration, rather than survival from mortality alone.

The residual analysis of the Hazard model showed more birds than expected on the open hunted sites, which we interpret as local compensation through net immigration or lower natural mortality (paragraph 2.3). Relative to closed sites, hunted sites held 3.4 % (median) more birds than predicted, but not consistently across pairs (A: 2.7 %, B: 8.9 %, C: −1.2 %). The net effect, combining the gain with the harvest loss relative to the closed sites, is shown in Table 3. At high effort the gain (3.7 %) offset only a small part of the harvest loss (11.6 %), leaving a net reduction of 8.5 % (95 % CrI 2.0–14.7 %) relative to closed sites, and full compensation was very unlikely (*P* (Δ_net_ ≥ 0) = 0.006). At low effort the gain (3.1 %) offset about half of the harvest loss (6.9 %), leaving a net reduction of 4.1 % (95 % CrI −1.6 to 9.6 %), and full compensation was unlikely (*P* (Δ_net_ ≥ 0) = 0.077).

**Table 3.**
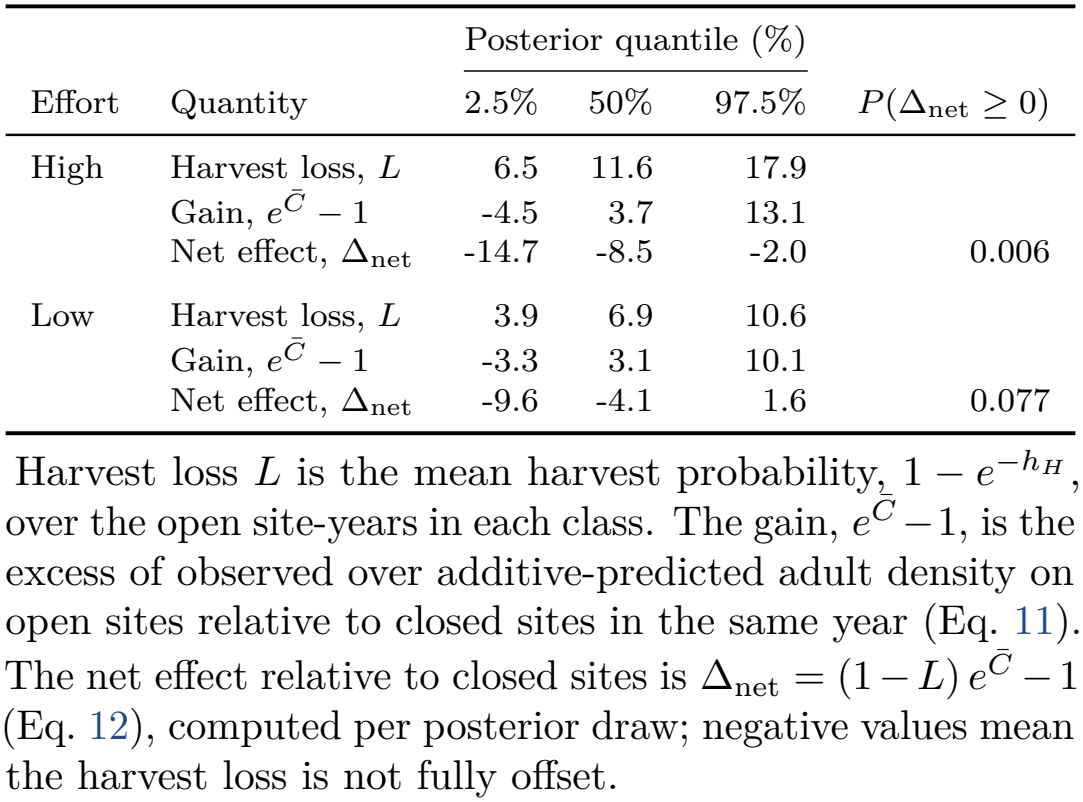
Harvest loss, compensatory gain and net effect of harvest on open (hunted) sites in site-years with high and low hunter effort, split at the center of the standardized effort scale (1.24 hd km^−2^; high: 14 site-years, median 2.44 hd km^−2^, range 1.44–6.57; low: 19 site-years, median 0.83, range 0.16–1.17). Values are posterior quantiles in percent; the last column gives the posterior probability that the gain fully offsets the harvest loss.

|  |  | Posterior quantile (%) |  |  |  |
|---|---|---|---|---|---|
| Effort | Quantity | 2.5% | 50% | 97.5% | *P*(Δ<sub>net</sub> ≥ 0) |
| High | Harvest loss, *L* | 6.5 | 11.6 | 17.9 |  |
|  | Gain, *e*<sup>*C̄*</sup> − 1 | -4.5 | 3.7 | 13.1 |  |
|  | Net effect, Δ<sub>net</sub> | -14.7 | -8.5 | -2.0 | 0.006 |
| Low | Harvest loss, *L* | 3.9 | 6.9 | 10.6 |  |
|  | Gain, *e*<sup>*C̄*</sup> − 1 | -3.3 | 3.1 | 10.1 |  |
|  | Net effect, Δ<sub>net</sub> | -9.6 | -4.1 | 1.6 | 0.077 |

Harvest loss *L* is the mean harvest probability, 1 − *e*<sup>−*h*<sub>*H*</sub></sup>, over the open site-years in each class. The gain, *e*<sup>*C̄*</sup> −1, is the excess of observed over additive-predicted adult density on open sites relative to closed sites in the same year (Eq. 11). The net effect relative to closed sites is Δ<sub>net</sub> = (1 − *L*) *e*<sup>*C̄*</sup> −1 (Eq. 12), computed per posterior draw; negative values mean the harvest loss is not fully offset.

**Table 4.** Total transect line length (km) surveyed per site and year in the August line-transect counts, 1996–2007, used as survey effort in the distance-sampling model. Letters denote site pairs; 0 = closed (unhunted), 1 = open (hunted). Site C0 was not surveyed in 1997 (–). Mean: mean over surveyed years.

| Site | 1996 | 1997 | 1998 | 1999 | 2000 | 2001 | 2002 | 2003 | 2004 | 2005 | 2006 | 2007 | Mean |
|---|---|---|---|---|---|---|---|---|---|---|---|---|---|
| A0 | 88.22 | 88.22 | 88.22 | 88.22 | 88.22 | 85.67 | 85.67 | 88.16 | 87.80 | 87.73 | 87.91 | 85.03 | 87.4 |
| A1 | 83.36 | 83.36 | 83.36 | 83.36 | 84.56 | 87.68 | 80.44 | 83.49 | 80.80 | 83.30 | 83.39 | 82.73 | 83.3 |
| B0 | 73.58 | 73.58 | 73.58 | 73.99 | 73.58 | 72.27 | 77.13 | 73.91 | 78.59 | 73.44 | 73.47 | 74.13 | 74.3 |
| B1 | 77.27 | 77.27 | 56.17 | 78.32 | 77.27 | 77.57 | 78.04 | 78.29 | 78.78 | 78.19 | 75.69 | 78.05 | 75.9 |
| C0 | 92.30 | – | 92.30 | 92.30 | 92.30 | 92.30 | 86.63 | 79.79 | 84.23 | 79.11 | 79.22 | 81.09 | 86.5 |
| C1 | 97.99 | 100.99 | 85.60 | 97.99 | 99.79 | 97.99 | 97.26 | 96.71 | 97.00 | 93.52 | 93.78 | 95.59 | 96.2 |

**Table 5.** Posterior predictive checks of the static distance-sampling model, by submodel. Obs is the statistic for the observed data (for the discrepancy statistics, its posterior median) and Rep its posterior median for replicated data; values of *P* (Rep *>* Obs) near 0 or 1 indicate lack of fit.

| Submodel | Test statistic | Obs | Rep | *P*(Rep > Obs) |
|---|---|---|---|---|
| Young per encounter | Mean young, 1 adult | 2.164 | 2.168 | 0.518 |
|  | Mean young, 2 adults | 4.017 | 4.000 | 0.447 |
|  | Mean young, ≥ 3 adults | 2.819 | 2.778 | 0.460 |
|  | P(young > 0), 1 adult | 0.504 | 0.504 | 0.496 |
|  | P(young > 0), 2 adults | 0.755 | 0.755 | 0.486 |
|  | P(young > 0), ≥ 3 adults | 0.375 | 0.375 | 0.487 |
|  | P(young ≥ 8), 1 adult | 0.081 | 0.065 | 0.012<sup>a</sup> |
|  | P(young ≥ 8), 2 adults | 0.181 | 0.166 | 0.100<sup>a</sup> |
|  | P(young ≥ 8), ≥ 3 adults | 0.171 | 0.157 | 0.288 |
| Encounter rate | Freeman–Tukey, site-years | 17.592 | 17.737 | 0.514 |
| Adult group size | χ<sup>2</sup>, size categories | 1.357 | 1.353 | 0.500 |
| Detection function | χ<sup>2</sup>, distance bins | 32.703 | 6.332 | 0.000<sup>b</sup> |

Young-count statistics are computed over all encounters, including those without young, by number of adults in the group. The last three rows are discrepancy statistics comparing counts with their model expectation.
<sup>a</sup> Borderline: very large broods are slightly under-predicted (see text).
<sup>b</sup> Lack of fit in the distance-bin counts, attributed to spatial clustering of encounters within transects (see text).

## 4. Discussion

Hunter effort measurably increased the mortality of willow ptarmigan, yet this cost was not detectable in year-to-year changes in abundance. The rise in harvest rate with effort, from a few percent at low effort to roughly thirty-five percent at the highest observed effort, is the firmest result of the study and the most important for management. The two models are consistent once their different questions are recognized: the Hazard model, constrained by the bag data, resolves how effort translates into harvest mortality, whereas the Gompertz model describes realized population growth, in which the harvest signal is small relative to variation in recruitment. Hunted sites held more birds than predicted from survival, which offset about half of the harvest loss at low effort, but only a third at high effort.

### Effort, harvest hazard and survival

The bridge to the population scale in the Hazard model is the relationship between effort, harvest rate and survival cost. Additivity is a structural assumption of our Hazard model: the natural and harvest hazards are added, so the corresponding survival probabilities multiply. Additive hazards therefore do not reduce survival by the full harvest rate; the reduction equals the harvest probability multiplied by survival in the absence of harvest, and this should not be mistaken for partial compensatory mortality. The high year-varying natural hazard is in accordance with the results on cause-specific mortality from the radio-collar study on willow ptarmigan in the region (Smith and Willebrand, 1999). That study concluded that harvest mortality was additive to natural mortality, and additive mortality has also been proposed in other studies using radio-collared grouse (Small, Holzwart, et al. 1991; Frye et al. 2022; Sandercock et al. 2011; Pedersen et al. 2004). The model cannot separate true mortality from net emigration, and the natural hazard is a net-loss hazard. This means that part of the estimated mortality can be the result of a redistribution of birds within the wider region. A favorable source–sink system of a small hunted fraction embedded in a large interior with low effort would buffer the regional population (Willebrand and Hörnell, 2001).

### Compensation on hunted sites

The residual analysis suggests local compensation: hunted sites held more birds than additive mortality predicts, offsetting about half of the harvest loss at low effort and a third at high effort, but not in all three pairs. A positive contrast can arise from net immigration or from lower natural mortality on hunted sites, which the shared natural hazard cannot separate. Shrinkage of the latent densities pulls the contrasts towards zero, and adjacent pair members may exchange birds, although radio-collared birds in the region gave no sign that closed sites supplied open ones (Smith and Willebrand, 1999).

### Breeding success dominates, and why the cost is not visible in abundance

The dominant driver of whether the population declines in a given year is reproductive success. An effect of productivity of about one (*γ* = 1.18, 95 % CrI 0.63–1.80) means that a change in autumn population size from breeding carries over about proportionally to the following spring (the interval includes one). Over-winter losses did not detectably offset good or poor breeding years within the same year. The productivity index, 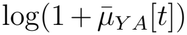, ranged from 0.88 (1996) to 1.18 (2003). Multiplied by *γ*, that gap is about 0.36 log units, so next year’s adult density is roughly 44 % higher after the best breeding year than after the poorest, all else equal. For contrast, one standard deviation of effort shifts next year’s density by about 2 %. Modelling brood occurrence and brood size separately helped recover this signal (Appendix A2). A strong effect of productivity was also found for the willow ptarmigan population in the Lövhögen area about 120–200 km south of the study sites (Willebrand, Hörnell-Willebrand, et al. 2025).

Two processes operating at different scales are likely to contribute to the rapid return towards site levels and to the buffering of the harvest signal. Every year, female natal dispersal redistributes birds across sites of varying harvest rates (Willebrand and Hörnell, 2001; Hörnell-Willebrand, Willebrand, et al. 2014). At the multiyear and regional scale, numerical and functional responses of a generalist predator community, with time lags, can stabilize total abundance and generate the synchronous fluctuations observed (Tornberg, Lindén, et al. 2013; Small, Marcström, et al. 1993; Breisjøberget et al. 2018; Kurki et al. 1997; Nielsen, 1999). The first-order return tendency of the Gompertz model absorbs both and cannot separate them.

Abundance estimates measure the consequence of harvest mortality after the compensating processes have operated. The additive harvest signal is clean at the level of demographic rates, but its consequence for abundance is buffered by density-dependent net survival and year-to-year variation in reproductive success; additive individual mortality is necessary but not sufficient for a detectable population-level effect. This can result in an apparent contradiction where radio-marking studies find additive mortality while abundance surveys struggle to detect any harvest signal. Both can be correct simultaneously: the mechanism is real, and the abundance signal is buffered below the threshold of detection. This can explain why Smith and Willebrand (1999) were not able to detect an anticipated decline in the harvested population. Abundance-only inference therefore risks missing masked additive mortality, whereas radio-only inference, read as though decline scaled directly with harvest rate, would ignore compensation; combining the two avoids both errors.

Willow ptarmigan across this region of the Swedish tundra show large annual fluctuations in density and chick production, but no clear directional trend attributable to harvest (unpublished reports on the annual counts to the counties of northern Sweden, compiled by M. Hörnell-Willebrand). Together, the unresolved effort coefficient and the absence of a regional trend in the monitoring show that population-level consequences of harvest at these effort levels cannot be detected with abundance data. The data are consistent with no effect, but also with substantial reductions in equilibrium density: relative to no harvest, the 95 % CrI for the long-run change at the upper threshold of 3 hd km^−2^ spans −54 to +42 % (Appendix A4). At sufficiently high effort, harvest will reduce populations, but how this translates into a measurable change in abundance over time depends on the stabilizing dynamics that the Hazard model does not contain. Our results are not evidence that harvest is benign, but rather that abundance-based monitoring alone cannot detect the impacts of the harvest levels documented here.

### Hunter behaviour

Hunters may reduce effort when birds are scarce, which would lower the harvest rate in poor years. We found no such response: effort was unrelated to productivity. Hunter satisfaction depends on much more than the bag (Vaske et al. 1986; Wam et al. 2012; Asmyhr et al. 2013), so effort is likely to continue at similar intensity in good and poor years, and harvest gives little self-correction when numbers are low.

### Implications for management

That the population-level average effect is hard to detect does not exclude the possibility that a small average effect could carry disproportionate weight in the poor reproductive years that matter most for harvest-dependent declines. This argument motivates an external limit on effort and precautionary management even where the directly detectable population signal is absent, rather than reliance on the harvest correcting itself.

Because harvest rate rises predictably with effort, we conclude that these results support the earlier suggestion that threshold management of harvest effort within management sites can be used to set allowable harvest rates well before the hunting season starts, a system that is easy to communicate to agencies and NGOs. Harvest effort is a quantity a manager can easily observe and regulate. Effort limits are common in commercial fisheries, where their success depends on whether catchability increases over time, and for other grouse species, but are usually combined with bag limits or adjusted to annual population counts, as in Finland (Baudron et al. 2010; Arlinghaus et al. 2015; Metsähallitus, 2026). In Finnmark County, Norway, quotas are set based on annual counts and models that generate near-term forecasts in an adaptive system (Henden et al. 2020).

### Limitations

Because persistent differences in abundance between sites are absorbed by the site intercepts *α*[*s*], any sustained effect of harvest on density is confounded with site differences.

The static and dynamic components were fitted in stages, with detection summaries passed forward as informative priors rather than estimated jointly; this two-stage approach does not fully propagate detection uncertainty into the dynamic estimates, so the dynamic-model intervals are likely slightly optimistic.

The Hazard model assumes additive harvest mortality by construction and therefore cannot test additivity against compensatory mortality; the additive interpretation rests on external radio-collar evidence. Neither dynamic model allows natural loss to depend on density in the previous year. If a lagged predator response lowers natural mortality in the year after a high harvest, part of the harvest loss would be compensated with a delay, which neither the within-year residual contrast (Eq. 11) nor the long-run projection in Appendix A4.4 would capture. The net effects in Table 3 may therefore overstate the population effects of harvest. One-step residuals suggest, though not significantly, that the Hazard model under-captures the effort effect in high-effort years (Appendix A7), so the survival costs at high effort are, if anything, conservative. Finally, the inference rests on a single species in one region over twelve years, with three open–closed site pairs. The agreement between the two models and with independent monitoring supports the effort–mortality result, but it does not extend the result beyond this system, predator community, or range of observed effort.

Whether harvest is compensated through density-dependent natural mortality therefore remains the main open question. Twelve years cannot answer it.

### Conclusions

Hunter effort raises the per-bird harvest rate and lowers annual survival by 3–7 percentage points within the recommended range of 1–3 hunter-days km^−2^, a clear and well-identified mechanism; the population effect is small on average but not demonstrably absent. We find no detectable decline in our dynamic model, in the long-term line-transect counts of eight sites in Jämtland (preliminary analysis of unpublished results), or in the long-term Lövhögen series at a harvest rate of about 10 % (Aanes et al. 2002; Willebrand, Hörnell-Willebrand, et al. 2025).

We conclude that the previous effort recommendations remain relevant: harvest effort should be kept around one hunter-day km^−2^ and should not exceed three hunter-days km^−2^ (Eriksson et al. 2006). The recommended upper limit of three hunter-days km^−2^ corresponds to a harvest probability of about 18 %, and a harvest probability of 30 % would be reached at about 5.2 hunter-days km^−2^. Present effort in the region (median 1.97 hunter-days km^−2^ in 2024/25) is locally above this level, which argues for annual follow-up of effort and counts. The hunting system provides little behavioral self-correction in the poor years when the margin for replacement is thinnest. Thus, relying on abundance alone, one would conclude *no effect*, whereas the mechanism we have identified, combined with weak self-regulation, indicates a genuine if hard-to-measure hazard.

Hunted sites gained more birds than survival predicted. The estimated gain from local compensation was similar at low and high effort (3.1 % and 3.7 %), so its share of the harvest loss fell as effort increased, from about half at low effort to a third at high effort. Natal dispersal is the likely source, although lower natural mortality on hunted sites cannot be excluded. We propose that harvest statistics be reported each year together with the abundance estimates from the monitoring programme, a low-cost addition that can largely be automated.

Effort set each year from preseason productivity and abundance would allow a higher average off-take and more precaution under a changing climate, but may not be feasible with the resources available, and neither managers nor hunters have asked for more effort. A fixed limit anchored to monitored effort and bag is therefore not a crude substitute for adaptive management, but an adequate design for state-managed land: precaution is the right response when a population-level consequence cannot be ruled out because it cannot be resolved.

We have begun an extended analysis using a Gompertz model with covariates from different trophic levels and the complete time series of the transect counts. The aim is to answer the question: How does a fluctuating prey population, whose recruitment is a stochastic process driven by spring conditions and microtine rodents, return towards an average level?

## Supporting information

Supplementary Material: Hunter effort increases willow ptarmigan mortality, but the cost is not visible in abundance: evaluating effort thresholds for

## Acknowledgements

We are grateful for all the support from the county management offices in Jämtland. This study would not have been possible without the dedicated counting teams and their highly qualified pointing dogs. The study received financial support from the MISTRA-foundation, the Swedish Hunters Association and the University of Inland Norway.

## Use of generative AI

The authors used Claude (Anthropic; Claude Opus models, accessed via claude.ai, 2025– 2026) as an assistant during this work. Uses comprised: (i) development, debugging and review of R and JAGS code, including identification of numerical errors in post-processing scripts; (ii) checking internal consistency between methods, results and parameter estimates; (iii) advice on the structure of the discussion and editing of the author’s text for length and clarity; and (iv) searches for relevant literature on effort-based harvest regulation. All analyses were specified, run and interpreted by the authors, all model code was checked and run by the author, and every cited source was verified against the original publication. The author takes full responsibility for the content.

## Data and script availability

Data and scripts are available online at https://doi.org/10.5281/zenodo.2211244.

## A. Model specifications and validation details

Priors were chosen to be weakly informative and biologically interpretable; where preliminary runs showed posteriors approaching prior boundaries, priors were refined to avoid artificial constraint of the posterior. MCMC settings were iteratively refined to ensure good convergence and effective sample size for all parameters; details are provided in Section A1. The static distance-sampling model is described in detail in Section A2, and the two dynamic models (Hazard and Gompertz) are described in Sections A3 and A4, respectively. Section A5 describes the standardization of hunter effort for prediction at the management thresholds, Section A6 the posterior predictive checks used to evaluate model fit, Section A7 a sensitivity check of the shared natural hazard, and Section A8 the full posterior parameter summaries.

### A1. MCMC settings

Three sets of MCMC settings were used during model development:

#### Initial evaluation

n.chains = 12, n.adapt = 300, n.burnin = 1 000, n.iter = 3 000, n.thin = 1. Used for early development and debugging.

#### Convergence checks

n.chains = 12, n.adapt = 500, n.burnin = 2 000, n.iter = 8 000, n.thin = 1. Used for intermediate versions close to convergence.

#### Final run

n.chains = 12, n.adapt = 500, n.burnin = 4 000, n.iter = 18 000, n.thin = 1. This yielded 168 000 posterior draws for inference.

The final run settings were chosen to ensure 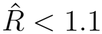 and effective sample size *>* 200 for all parameters, which are standard thresholds for MCMC convergence and adequate posterior sampling. All reported parameters had 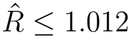 and *n*_eff_ ≥ 500; values of 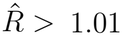, a stricter criterion, are flagged in Tables 1, 7 and 8. Non-centered parameterization (*z* ∼ N (0, 1); parameter = *µ* + *σ* · *z*) was used throughout to improve chain mixing, following standard advice for hierarchical models with small numbers of groups.

**Table 6.**
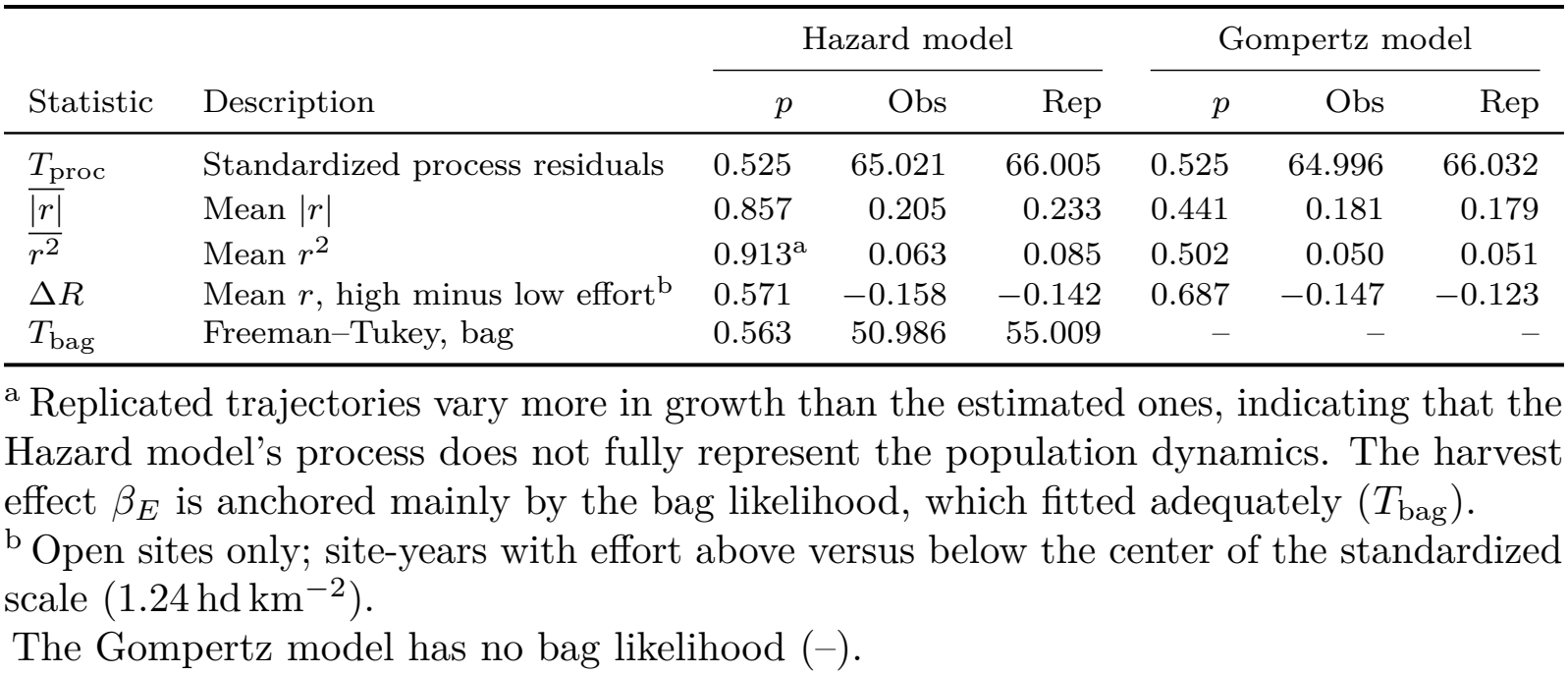
Posterior predictive checks of the Hazard and Gompertz state-space models. Obs is the posterior mean of each statistic for the estimated latent trajectories (for *T*_bag_, for the observed bags) and Rep its posterior mean for data replicated from the model; *p* = *P* (Rep *>* Obs), with values near 0 or 1 indicating lack of fit. Growth is the annual change in log adult density, *r* = log *D*_*A*_[*t* + 1] − log *D*_*A*_[*t*].

|  |  | Hazard model |  |  | Gompertz model |  |  |
|---|---|---|---|---|---|---|---|
| Statistic | Description | *p* | Obs | Rep | *p* | Obs | Rep |
| *T*<sub>proc</sub> | Standardized process residuals | 0.525 | 65.021 | 66.005 | 0.525 | 64.996 | 66.032 |
| $\overline{\lvert r\rvert}$ | Mean \|*r*\| | 0.857 | 0.205 | 0.233 | 0.441 | 0.181 | 0.179 |
| $\overline{r^2}$ | Mean *r*<sup>2</sup> | 0.913<sup>a</sup> | 0.063 | 0.085 | 0.502 | 0.050 | 0.051 |
| Δ*R* | Mean *r*, high minus low effort<sup>b</sup> | 0.571 | −0.158 | −0.142 | 0.687 | −0.147 | −0.123 |
| *T*<sub>bag</sub> | Freeman–Tukey, bag | 0.563 | 50.986 | 55.009 | – | – | – |

<sup>a</sup> Replicated trajectories vary more in growth than the estimated ones, indicating that the Hazard model's process does not fully represent the population dynamics. The harvest effect *β*<sub>*E*</sub> is anchored mainly by the bag likelihood, which fitted adequately (*T*<sub>bag</sub>).
<sup>b</sup> Open sites only; site-years with effort above versus below the center of the standardized scale (1.24 hd km<sup>−2</sup>).
The Gompertz model has no bag likelihood (–).

**Table 7.**
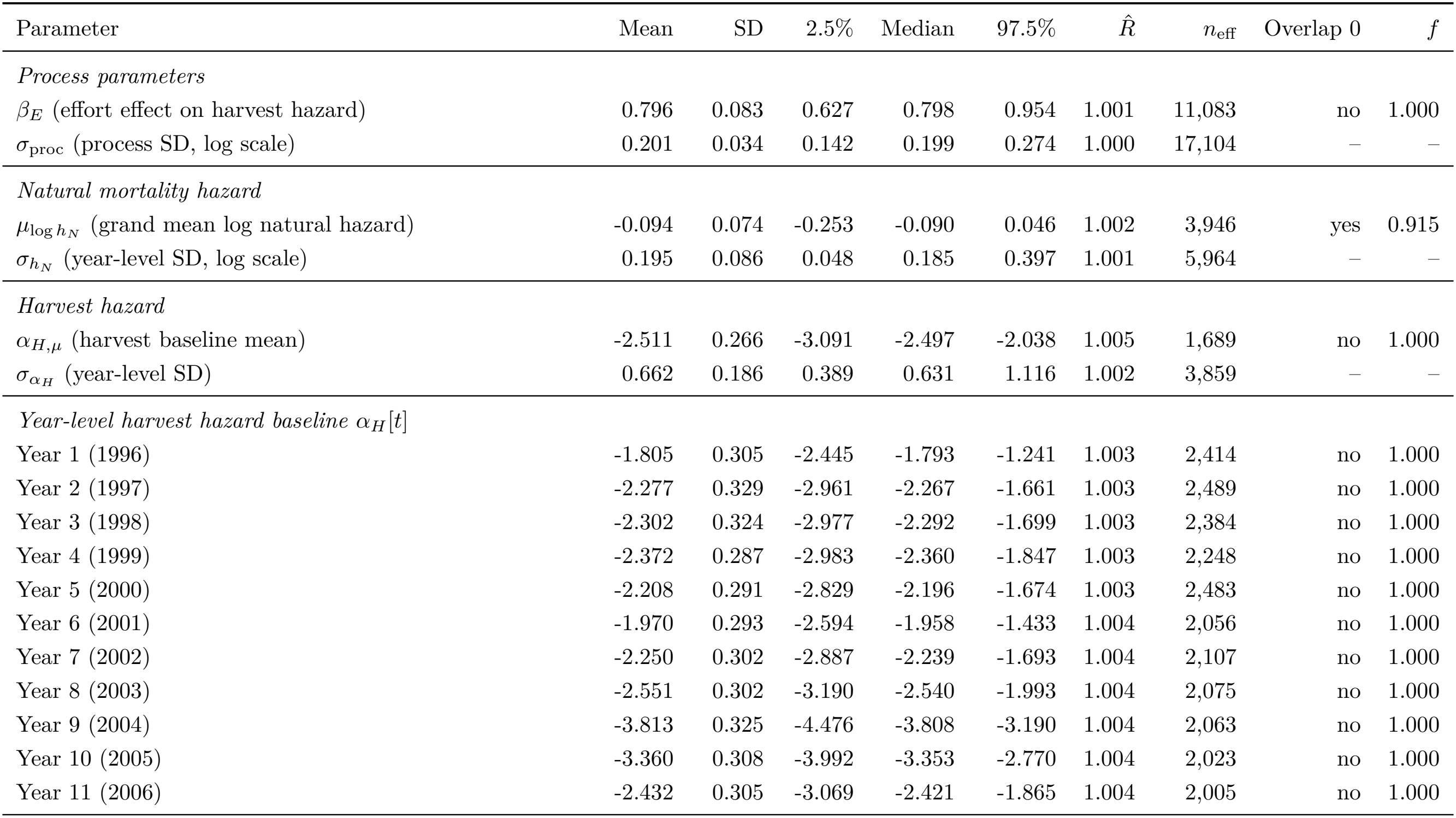
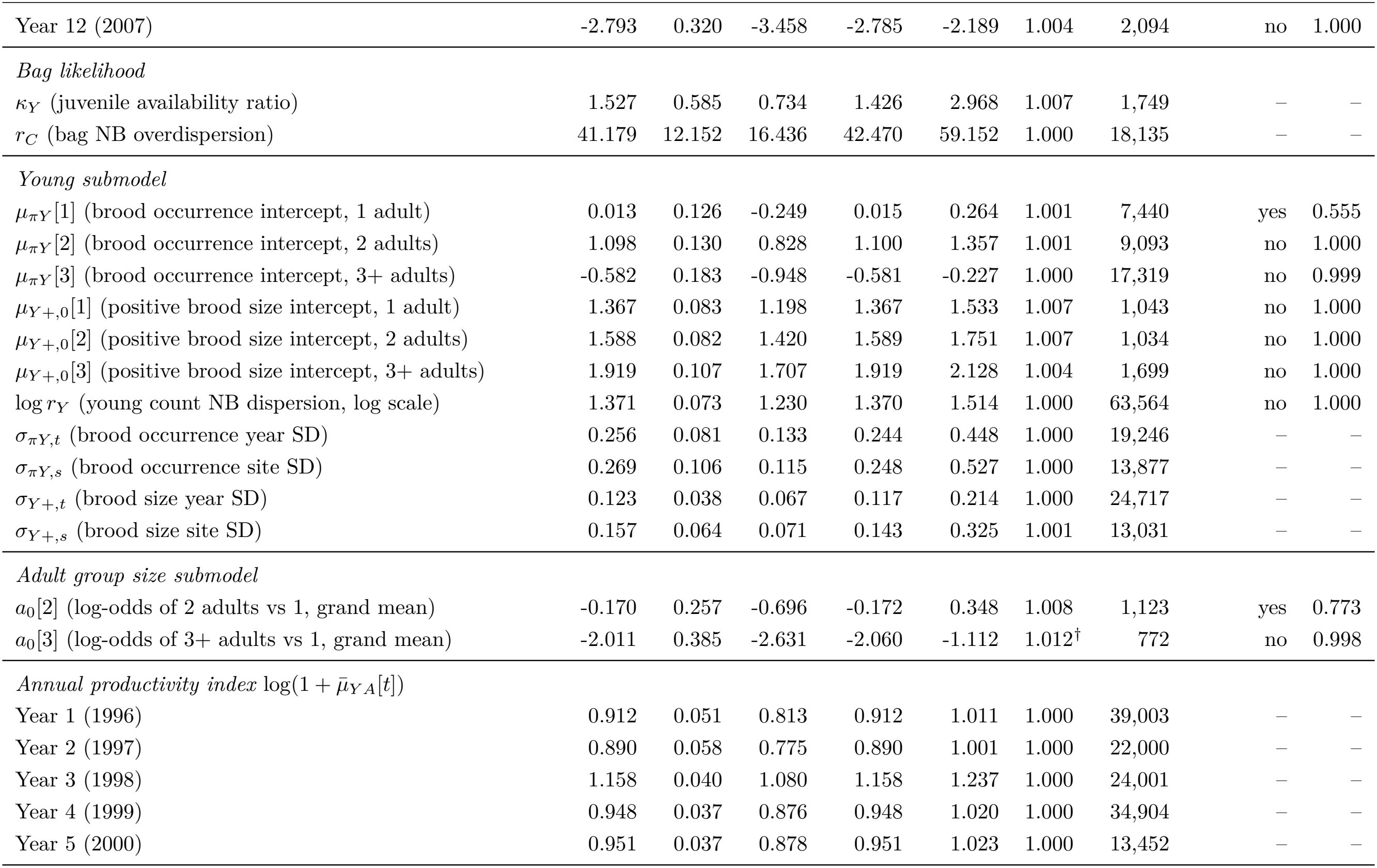
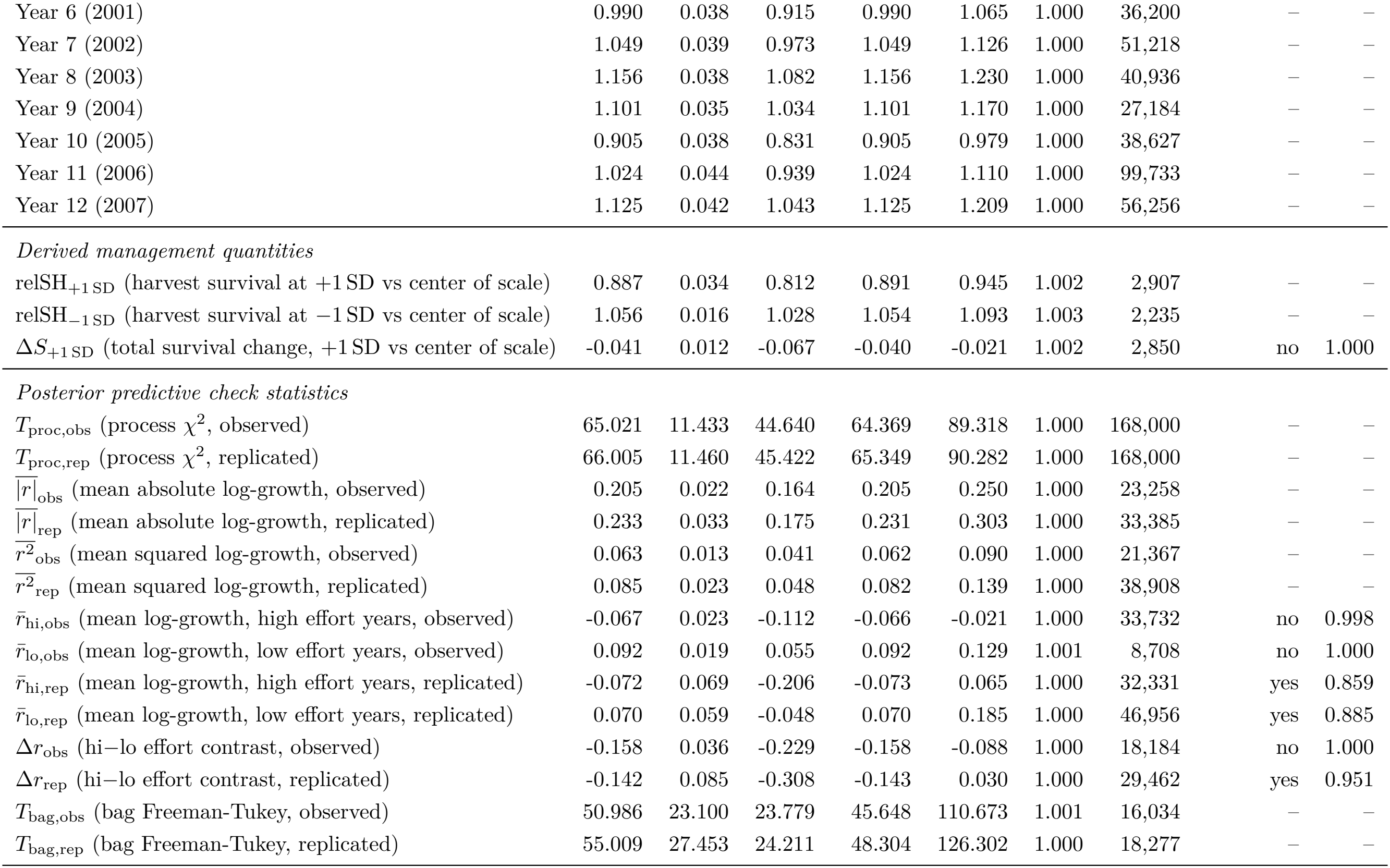
Full posterior summary for the Hazard state-space model, based on 168,000 posterior draws from 12 chains. Rows are grouped by submodel and include derived management quantities and posterior predictive check statistics. Columns give the posterior mean, SD, 2.5 % quantile, median and 97.5 % quantile; 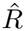, Gelman–Rubin statistic 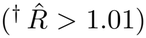; *n*_eff_, effective sample size; Overlap 0, whether the 95 % credible interval includes zero; *f*, proportion of the posterior with the same sign as the mean. Overlap 0 and *f* are not shown (–) for quantities that are positive by definition. Derived survival quantities are relative to the center of the standardized effort scale (1.24 hd km^−2^).

| Parameter | Mean | SD | 2.5% | Median | 97.5% | *R̂* | *n*<sub>eff</sub> | Overlap 0 | *f* |
|---|---|---|---|---|---|---|---|---|---|
| *Process parameters* |  |  |  |  |  |  |  |  |  |
| β<sub>*E*</sub> (effort effect on harvest hazard) | 0.796 | 0.083 | 0.627 | 0.798 | 0.954 | 1.001 | 11,083 | no | 1.000 |
| σ<sub>proc</sub> (process SD, log scale) | 0.201 | 0.034 | 0.142 | 0.199 | 0.274 | 1.000 | 17,104 | – | – |
| *Natural mortality hazard* |  |  |  |  |  |  |  |  |  |
| μ<sub>log *h*<sub>*N*</sub></sub> (grand mean log natural hazard) | -0.094 | 0.074 | -0.253 | -0.090 | 0.046 | 1.002 | 3,946 | yes | 0.915 |
| σ<sub>*h*<sub>*N*</sub></sub> (year-level SD, log scale) | 0.195 | 0.086 | 0.048 | 0.185 | 0.397 | 1.001 | 5,964 | – | – |
| *Harvest hazard* |  |  |  |  |  |  |  |  |  |
| α<sub>*H*,μ</sub> (harvest baseline mean) | -2.511 | 0.266 | -3.091 | -2.497 | -2.038 | 1.005 | 1,689 | no | 1.000 |
| σ<sub>α<sub>*H*</sub></sub> (year-level SD) | 0.662 | 0.186 | 0.389 | 0.631 | 1.116 | 1.002 | 3,859 | – | – |
| *Year-level harvest hazard baseline* α<sub>*H*</sub>[*t*] |  |  |  |  |  |  |  |  |  |
| Year 1 (1996) | -1.805 | 0.305 | -2.445 | -1.793 | -1.241 | 1.003 | 2,414 | no | 1.000 |
| Year 2 (1997) | -2.277 | 0.329 | -2.961 | -2.267 | -1.661 | 1.003 | 2,489 | no | 1.000 |
| Year 3 (1998) | -2.302 | 0.324 | -2.977 | -2.292 | -1.699 | 1.003 | 2,384 | no | 1.000 |
| Year 4 (1999) | -2.372 | 0.287 | -2.983 | -2.360 | -1.847 | 1.003 | 2,248 | no | 1.000 |
| Year 5 (2000) | -2.208 | 0.291 | -2.829 | -2.196 | -1.674 | 1.003 | 2,483 | no | 1.000 |
| Year 6 (2001) | -1.970 | 0.293 | -2.594 | -1.958 | -1.433 | 1.004 | 2,056 | no | 1.000 |
| Year 7 (2002) | -2.250 | 0.302 | -2.887 | -2.239 | -1.693 | 1.004 | 2,107 | no | 1.000 |
| Year 8 (2003) | -2.551 | 0.302 | -3.190 | -2.540 | -1.993 | 1.004 | 2,075 | no | 1.000 |
| Year 9 (2004) | -3.813 | 0.325 | -4.476 | -3.808 | -3.190 | 1.004 | 2,063 | no | 1.000 |
| Year 10 (2005) | -3.360 | 0.308 | -3.992 | -3.353 | -2.770 | 1.004 | 2,023 | no | 1.000 |
| Year 11 (2006) | -2.432 | 0.305 | -3.069 | -2.421 | -1.865 | 1.004 | 2,005 | no | 1.000 |
| Year 12 (2007) | -2.793 | 0.320 | -3.458 | -2.785 | -2.189 | 1.004 | 2,094 | no | 1.000 |
| *Bag likelihood* |  |  |  |  |  |  |  |  |  |
| κ<sub>*Y*</sub> (juvenile availability ratio) | 1.527 | 0.585 | 0.734 | 1.426 | 2.968 | 1.007 | 1,749 | – | – |
| *r*<sub>*C*</sub> (bag NB overdispersion) | 41.179 | 12.152 | 16.436 | 42.470 | 59.152 | 1.000 | 18,135 | – | – |
| *Young submodel* |  |  |  |  |  |  |  |  |  |
| μ<sub>π*Y*</sub>[1] (brood occurrence intercept, 1 adult) | 0.013 | 0.126 | -0.249 | 0.015 | 0.264 | 1.001 | 7,440 | yes | 0.555 |
| μ<sub>π*Y*</sub>[2] (brood occurrence intercept, 2 adults) | 1.098 | 0.130 | 0.828 | 1.100 | 1.357 | 1.001 | 9,093 | no | 1.000 |
| μ<sub>π*Y*</sub>[3] (brood occurrence intercept, 3+ adults) | -0.582 | 0.183 | -0.948 | -0.581 | -0.227 | 1.000 | 17,319 | no | 0.999 |
| μ<sub>*Y*+,0</sub>[1] (positive brood size intercept, 1 adult) | 1.367 | 0.083 | 1.198 | 1.367 | 1.533 | 1.007 | 1,043 | no | 1.000 |
| μ<sub>*Y*+,0</sub>[2] (positive brood size intercept, 2 adults) | 1.588 | 0.082 | 1.420 | 1.589 | 1.751 | 1.007 | 1,034 | no | 1.000 |
| μ<sub>*Y*+,0</sub>[3] (positive brood size intercept, 3+ adults) | 1.919 | 0.107 | 1.707 | 1.919 | 2.128 | 1.004 | 1,699 | no | 1.000 |
| log *r*<sub>*Y*</sub> (young count NB dispersion, log scale) | 1.371 | 0.073 | 1.230 | 1.370 | 1.514 | 1.000 | 63,564 | no | 1.000 |
| σ<sub>π*Y*,*t*</sub> (brood occurrence year SD) | 0.256 | 0.081 | 0.133 | 0.244 | 0.448 | 1.000 | 19,246 | – | – |
| σ<sub>π*Y*,*s*</sub> (brood occurrence site SD) | 0.269 | 0.106 | 0.115 | 0.248 | 0.527 | 1.000 | 13,877 | – | – |
| σ<sub>*Y*+,*t*</sub> (brood size year SD) | 0.123 | 0.038 | 0.067 | 0.117 | 0.214 | 1.000 | 24,717 | – | – |
| σ<sub>*Y*+,*s*</sub> (brood size site SD) | 0.157 | 0.064 | 0.071 | 0.143 | 0.325 | 1.001 | 13,031 | – | – |
| *Adult group size submodel* |  |  |  |  |  |  |  |  |  |
| *a*<sub>0</sub>[2] (log-odds of 2 adults vs 1, grand mean) | -0.170 | 0.257 | -0.696 | -0.172 | 0.348 | 1.008 | 1,123 | yes | 0.773 |
| *a*<sub>0</sub>[3] (log-odds of 3+ adults vs 1, grand mean) | -2.011 | 0.385 | -2.631 | -2.060 | -1.112 | 1.012<sup>†</sup> | 772 | no | 0.998 |
| *Annual productivity index* log(1 + μ̄<sub>*YA*</sub>[*t*]) |  |  |  |  |  |  |  |  |  |
| Year 1 (1996) | 0.912 | 0.051 | 0.813 | 0.912 | 1.011 | 1.000 | 39,003 | – | – |
| Year 2 (1997) | 0.890 | 0.058 | 0.775 | 0.890 | 1.001 | 1.000 | 22,000 | – | – |
| Year 3 (1998) | 1.158 | 0.040 | 1.080 | 1.158 | 1.237 | 1.000 | 24,001 | – | – |
| Year 4 (1999) | 0.948 | 0.037 | 0.876 | 0.948 | 1.020 | 1.000 | 34,904 | – | – |
| Year 5 (2000) | 0.951 | 0.037 | 0.878 | 0.951 | 1.023 | 1.000 | 13,452 | – | – |
| Year 6 (2001) | 0.990 | 0.038 | 0.915 | 0.990 | 1.065 | 1.000 | 36,200 | – | – |
| Year 7 (2002) | 1.049 | 0.039 | 0.973 | 1.049 | 1.126 | 1.000 | 51,218 | – | – |
| Year 8 (2003) | 1.156 | 0.038 | 1.082 | 1.156 | 1.230 | 1.000 | 40,936 | – | – |
| Year 9 (2004) | 1.101 | 0.035 | 1.034 | 1.101 | 1.170 | 1.000 | 27,184 | – | – |
| Year 10 (2005) | 0.905 | 0.038 | 0.831 | 0.905 | 0.979 | 1.000 | 38,627 | – | – |
| Year 11 (2006) | 1.024 | 0.044 | 0.939 | 1.024 | 1.110 | 1.000 | 99,733 | – | – |
| Year 12 (2007) | 1.125 | 0.042 | 1.043 | 1.125 | 1.209 | 1.000 | 56,256 | – | – |
| *Derived management quantities* |  |  |  |  |  |  |  |  |  |
| relSH<sub>+1 SD</sub> (harvest survival at +1 SD vs center of scale) | 0.887 | 0.034 | 0.812 | 0.891 | 0.945 | 1.002 | 2,907 | – | – |
| relSH<sub>−1 SD</sub> (harvest survival at −1 SD vs center of scale) | 1.056 | 0.016 | 1.028 | 1.054 | 1.093 | 1.003 | 2,235 | – | – |
| Δ*S*<sub>+1 SD</sub> (total survival change, +1 SD vs center of scale) | -0.041 | 0.012 | -0.067 | -0.040 | -0.021 | 1.002 | 2,850 | no | 1.000 |
| *Posterior predictive check statistics* |  |  |  |  |  |  |  |  |  |
| *T*<sub>proc,obs</sub> (process χ<sup>2</sup>, observed) | 65.021 | 11.433 | 44.640 | 64.369 | 89.318 | 1.000 | 168,000 | – | – |
| *T*<sub>proc,rep</sub> (process χ<sup>2</sup>, replicated) | 66.005 | 11.460 | 45.422 | 65.349 | 90.282 | 1.000 | 168,000 | – | – |
| $\overline{\lvert r\rvert}$<sub>obs</sub> (mean absolute log-growth, observed) | 0.205 | 0.022 | 0.164 | 0.205 | 0.250 | 1.000 | 23,258 | – | – |
| $\overline{\lvert r\rvert}$<sub>rep</sub> (mean absolute log-growth, replicated) | 0.233 | 0.033 | 0.175 | 0.231 | 0.303 | 1.000 | 33,385 | – | – |
| $\overline{r^2}$<sub>obs</sub> (mean squared log-growth, observed) | 0.063 | 0.013 | 0.041 | 0.062 | 0.090 | 1.000 | 21,367 | – | – |
| $\overline{r^2}$<sub>rep</sub> (mean squared log-growth, replicated) | 0.085 | 0.023 | 0.048 | 0.082 | 0.139 | 1.000 | 38,908 | – | – |
| *r̄*<sub>hi,obs</sub> (mean log-growth, high effort years, observed) | -0.067 | 0.023 | -0.112 | -0.066 | -0.021 | 1.000 | 33,732 | no | 0.998 |
| *r̄*<sub>lo,obs</sub> (mean log-growth, low effort years, observed) | 0.092 | 0.019 | 0.055 | 0.092 | 0.129 | 1.001 | 8,708 | no | 1.000 |
| *r̄*<sub>hi,rep</sub> (mean log-growth, high effort years, replicated) | -0.072 | 0.069 | -0.206 | -0.073 | 0.065 | 1.000 | 32,331 | yes | 0.859 |
| *r̄*<sub>lo,rep</sub> (mean log-growth, low effort years, replicated) | 0.070 | 0.059 | -0.048 | 0.070 | 0.185 | 1.000 | 46,956 | yes | 0.885 |
| Δ*r*<sub>obs</sub> (hi−lo effort contrast, observed) | -0.158 | 0.036 | -0.229 | -0.158 | -0.088 | 1.000 | 18,184 | no | 1.000 |
| Δ*r*<sub>rep</sub> (hi−lo effort contrast, replicated) | -0.142 | 0.085 | -0.308 | -0.143 | 0.030 | 1.000 | 29,462 | yes | 0.951 |
| *T*<sub>bag,obs</sub> (bag Freeman-Tukey, observed) | 50.986 | 23.100 | 23.779 | 45.648 | 110.673 | 1.001 | 16,034 | – | – |
| *T*<sub>bag,rep</sub> (bag Freeman-Tukey, replicated) | 55.009 | 27.453 | 24.211 | 48.304 | 126.302 | 1.000 | 18,277 | – | – |

**Table 8.**
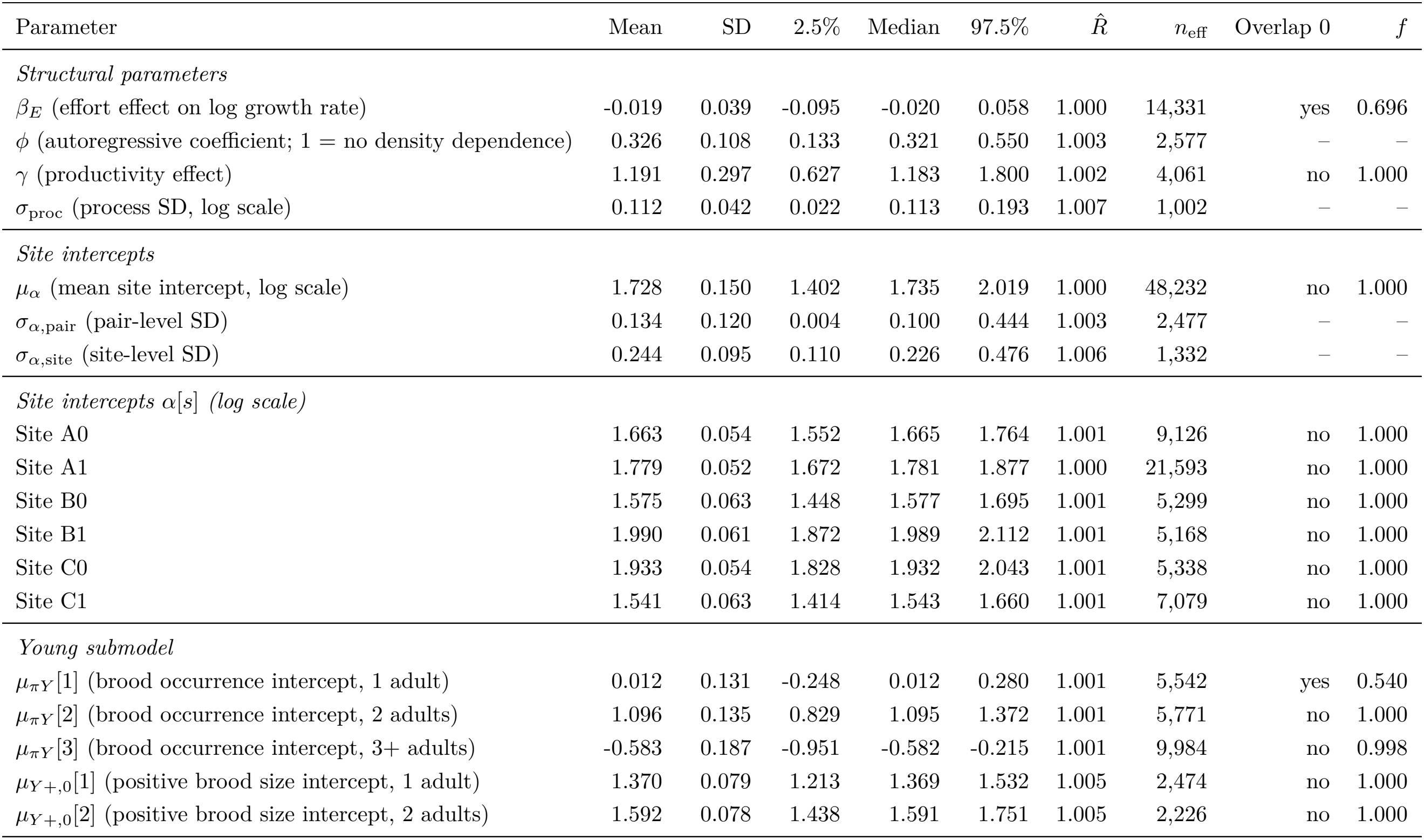
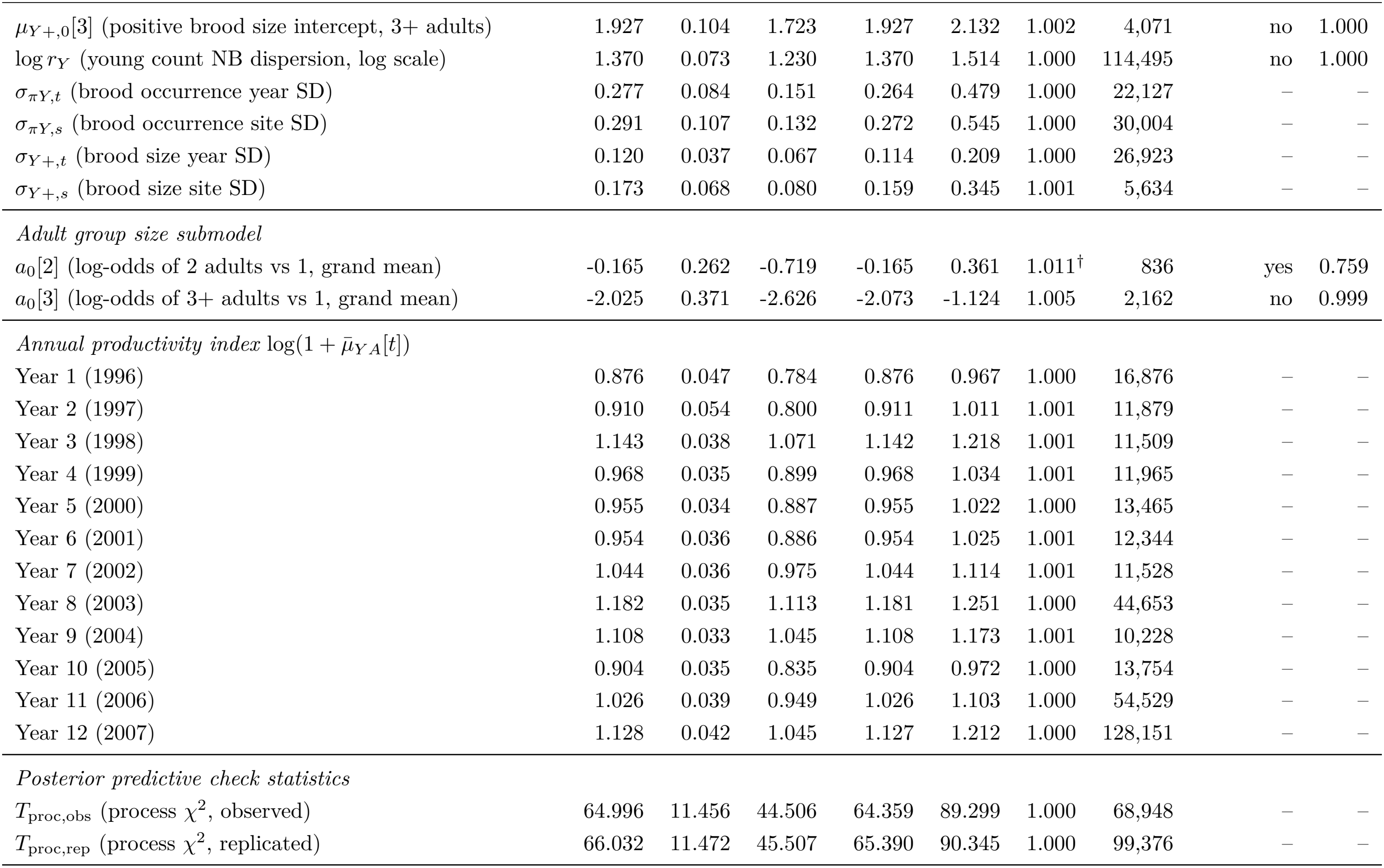
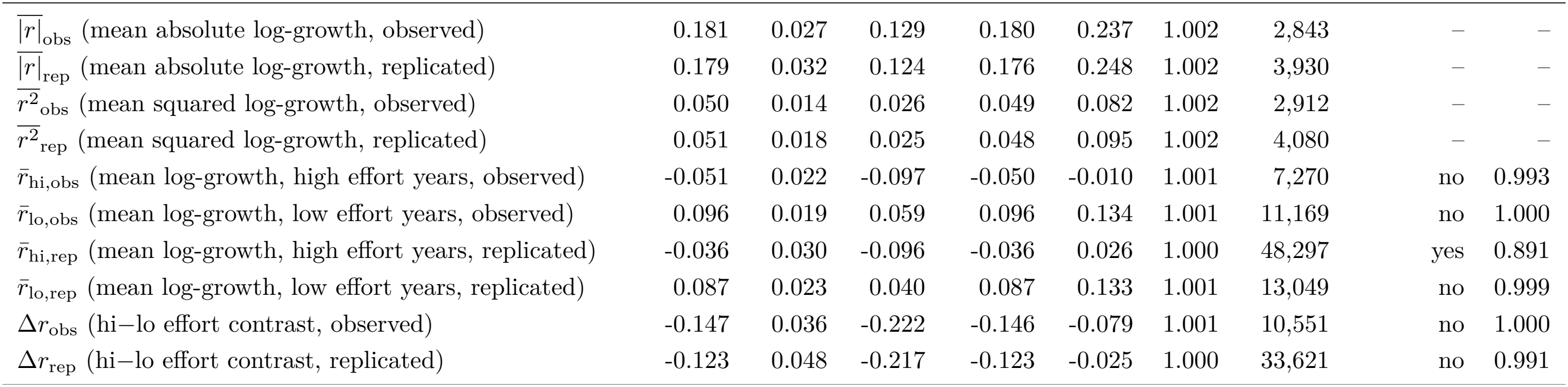
Full posterior summary for the Gompertz state-space model (Eq. 6), based on 168,000 posterior draws from 12 chains. *ϕ* = 1 means no density dependence and 1 − *ϕ* is the proportion of a deviation from the site’s equilibrium level removed per year. Columns and symbols as in Table 7.

| Parameter | Mean | SD | 2.5% | Median | 97.5% | R̂ | n<sub>eff</sub> | Overlap 0 | f |
|---|---|---|---|---|---|---|---|---|---|
| *Structural parameters* |  |  |  |  |  |  |  |  |  |
| β<sub>E</sub> (effort effect on log growth rate) | -0.019 | 0.039 | -0.095 | -0.020 | 0.058 | 1.000 | 14,331 | yes | 0.696 |
| ϕ (autoregressive coefficient; 1 = no density dependence) | 0.326 | 0.108 | 0.133 | 0.321 | 0.550 | 1.003 | 2,577 | – | – |
| γ (productivity effect) | 1.191 | 0.297 | 0.627 | 1.183 | 1.800 | 1.002 | 4,061 | no | 1.000 |
| σ<sub>proc</sub> (process SD, log scale) | 0.112 | 0.042 | 0.022 | 0.113 | 0.193 | 1.007 | 1,002 | – | – |
| *Site intercepts* |  |  |  |  |  |  |  |  |  |
| μ<sub>α</sub> (mean site intercept, log scale) | 1.728 | 0.150 | 1.402 | 1.735 | 2.019 | 1.000 | 48,232 | no | 1.000 |
| σ<sub>α,pair</sub> (pair-level SD) | 0.134 | 0.120 | 0.004 | 0.100 | 0.444 | 1.003 | 2,477 | – | – |
| σ<sub>α,site</sub> (site-level SD) | 0.244 | 0.095 | 0.110 | 0.226 | 0.476 | 1.006 | 1,332 | – | – |
| *Site intercepts* α[s] *(log scale)* |  |  |  |  |  |  |  |  |  |
| Site A0 | 1.663 | 0.054 | 1.552 | 1.665 | 1.764 | 1.001 | 9,126 | no | 1.000 |
| Site A1 | 1.779 | 0.052 | 1.672 | 1.781 | 1.877 | 1.000 | 21,593 | no | 1.000 |
| Site B0 | 1.575 | 0.063 | 1.448 | 1.577 | 1.695 | 1.001 | 5,299 | no | 1.000 |
| Site B1 | 1.990 | 0.061 | 1.872 | 1.989 | 2.112 | 1.001 | 5,168 | no | 1.000 |
| Site C0 | 1.933 | 0.054 | 1.828 | 1.932 | 2.043 | 1.001 | 5,338 | no | 1.000 |
| Site C1 | 1.541 | 0.063 | 1.414 | 1.543 | 1.660 | 1.001 | 7,079 | no | 1.000 |
| *Young submodel* |  |  |  |  |  |  |  |  |  |
| μ<sub>πY</sub>[1] (brood occurrence intercept, 1 adult) | 0.012 | 0.131 | -0.248 | 0.012 | 0.280 | 1.001 | 5,542 | yes | 0.540 |
| μ<sub>πY</sub>[2] (brood occurrence intercept, 2 adults) | 1.096 | 0.135 | 0.829 | 1.095 | 1.372 | 1.001 | 5,771 | no | 1.000 |
| μ<sub>πY</sub>[3] (brood occurrence intercept, 3+ adults) | -0.583 | 0.187 | -0.951 | -0.582 | -0.215 | 1.001 | 9,984 | no | 0.998 |
| μ<sub>Y+,0</sub>[1] (positive brood size intercept, 1 adult) | 1.370 | 0.079 | 1.213 | 1.369 | 1.532 | 1.005 | 2,474 | no | 1.000 |
| μ<sub>Y+,0</sub>[2] (positive brood size intercept, 2 adults) | 1.592 | 0.078 | 1.438 | 1.591 | 1.751 | 1.005 | 2,226 | no | 1.000 |
| μ<sub>Y+,0</sub>[3] (positive brood size intercept, 3+ adults) | 1.927 | 0.104 | 1.723 | 1.927 | 2.132 | 1.002 | 4,071 | no | 1.000 |
| log r<sub>Y</sub> (young count NB dispersion, log scale) | 1.370 | 0.073 | 1.230 | 1.370 | 1.514 | 1.000 | 114,495 | no | 1.000 |
| σ<sub>πY,t</sub> (brood occurrence year SD) | 0.277 | 0.084 | 0.151 | 0.264 | 0.479 | 1.000 | 22,127 | – | – |
| σ<sub>πY,s</sub> (brood occurrence site SD) | 0.291 | 0.107 | 0.132 | 0.272 | 0.545 | 1.000 | 30,004 | – | – |
| σ<sub>Y+,t</sub> (brood size year SD) | 0.120 | 0.037 | 0.067 | 0.114 | 0.209 | 1.000 | 26,923 | – | – |
| σ<sub>Y+,s</sub> (brood size site SD) | 0.173 | 0.068 | 0.080 | 0.159 | 0.345 | 1.001 | 5,634 | – | – |
| *Adult group size submodel* |  |  |  |  |  |  |  |  |  |
| a<sub>0</sub>[2] (log-odds of 2 adults vs 1, grand mean) | -0.165 | 0.262 | -0.719 | -0.165 | 0.361 | 1.011<sup>†</sup> | 836 | yes | 0.759 |
| a<sub>0</sub>[3] (log-odds of 3+ adults vs 1, grand mean) | -2.025 | 0.371 | -2.626 | -2.073 | -1.124 | 1.005 | 2,162 | no | 0.999 |
| *Annual productivity index* log(1 + μ̄<sub>YA</sub>[t]) |  |  |  |  |  |  |  |  |  |
| Year 1 (1996) | 0.876 | 0.047 | 0.784 | 0.876 | 0.967 | 1.000 | 16,876 | – | – |
| Year 2 (1997) | 0.910 | 0.054 | 0.800 | 0.911 | 1.011 | 1.001 | 11,879 | – | – |
| Year 3 (1998) | 1.143 | 0.038 | 1.071 | 1.142 | 1.218 | 1.001 | 11,509 | – | – |
| Year 4 (1999) | 0.968 | 0.035 | 0.899 | 0.968 | 1.034 | 1.001 | 11,965 | – | – |
| Year 5 (2000) | 0.955 | 0.034 | 0.887 | 0.955 | 1.022 | 1.000 | 13,465 | – | – |
| Year 6 (2001) | 0.954 | 0.036 | 0.886 | 0.954 | 1.025 | 1.001 | 12,344 | – | – |
| Year 7 (2002) | 1.044 | 0.036 | 0.975 | 1.044 | 1.114 | 1.001 | 11,528 | – | – |
| Year 8 (2003) | 1.182 | 0.035 | 1.113 | 1.181 | 1.251 | 1.000 | 44,653 | – | – |
| Year 9 (2004) | 1.108 | 0.033 | 1.045 | 1.108 | 1.173 | 1.001 | 10,228 | – | – |
| Year 10 (2005) | 0.904 | 0.035 | 0.835 | 0.904 | 0.972 | 1.000 | 13,754 | – | – |
| Year 11 (2006) | 1.026 | 0.039 | 0.949 | 1.026 | 1.103 | 1.000 | 54,529 | – | – |
| Year 12 (2007) | 1.128 | 0.042 | 1.045 | 1.127 | 1.212 | 1.000 | 128,151 | – | – |
| *Posterior predictive check statistics* |  |  |  |  |  |  |  |  |  |
| T<sub>proc,obs</sub> (process χ<sup>2</sup>, observed) | 64.996 | 11.456 | 44.506 | 64.359 | 89.299 | 1.000 | 68,948 | – | – |
| T<sub>proc,rep</sub> (process χ<sup>2</sup>, replicated) | 66.032 | 11.472 | 45.507 | 65.390 | 90.345 | 1.000 | 99,376 | – | – |
| $\overline{\|r\|}_{\mathrm{obs}}$ (mean absolute log-growth, observed) | 0.181 | 0.027 | 0.129 | 0.180 | 0.237 | 1.002 | 2,843 | – | – |
| $\overline{\|r\|}_{\mathrm{rep}}$ (mean absolute log-growth, replicated) | 0.179 | 0.032 | 0.124 | 0.176 | 0.248 | 1.002 | 3,930 | – | – |
| $\overline{r^2}_{\mathrm{obs}}$ (mean squared log-growth, observed) | 0.050 | 0.014 | 0.026 | 0.049 | 0.082 | 1.002 | 2,912 | – | – |
| $\overline{r^2}_{\mathrm{rep}}$ (mean squared log-growth, replicated) | 0.051 | 0.018 | 0.025 | 0.048 | 0.095 | 1.002 | 4,080 | – | – |
| r̄<sub>hi,obs</sub> (mean log-growth, high effort years, observed) | -0.051 | 0.022 | -0.097 | -0.050 | -0.010 | 1.001 | 7,270 | no | 0.993 |
| r̄<sub>lo,obs</sub> (mean log-growth, low effort years, observed) | 0.096 | 0.019 | 0.059 | 0.096 | 0.134 | 1.001 | 11,169 | no | 1.000 |
| r̄<sub>hi,rep</sub> (mean log-growth, high effort years, replicated) | -0.036 | 0.030 | -0.096 | -0.036 | 0.026 | 1.000 | 48,297 | yes | 0.891 |
| r̄<sub>lo,rep</sub> (mean log-growth, low effort years, replicated) | 0.087 | 0.023 | 0.040 | 0.087 | 0.133 | 1.001 | 13,049 | no | 0.999 |
| Δr<sub>obs</sub> (hi−lo effort contrast, observed) | -0.147 | 0.036 | -0.222 | -0.146 | -0.079 | 1.001 | 10,551 | no | 1.000 |
| Δr<sub>rep</sub> (hi−lo effort contrast, replicated) | -0.123 | 0.048 | -0.217 | -0.123 | -0.025 | 1.000 | 33,621 | no | 0.991 |

### A2. Static distance-sampling model

#### A2.1 Detection function and shape parameters

The hazard-rate detection function is:

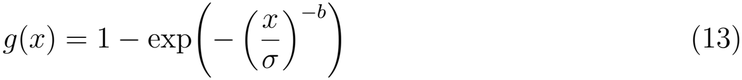

The shape parameter *b* was fixed rather than estimated freely, because freely estimating *b* creates a near-ridge in the posterior between *b* and *σ*, causing slow chain mixing without improving the fit to the quantities of scientific interest (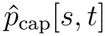 and *D*_grp_[*s, t*]). Fixed values were chosen from the posterior medians of runs on the corrected data in which *b* was freely estimated:

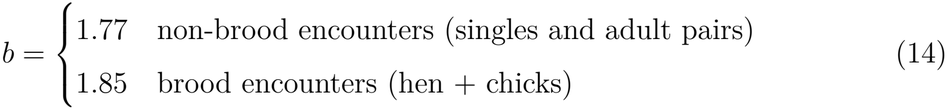

Fixing *b* does not preclude variation in the detection function across site-years, because *σ* retains a full hierarchical structure.

#### A2.2 Hierarchical scale parameter

The scale *σ* varies across sites, years, and encounter types:

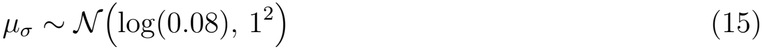

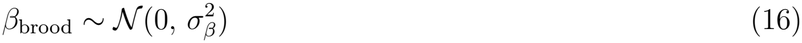

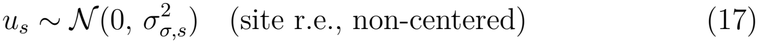

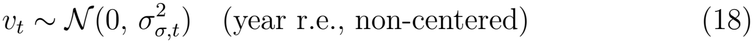

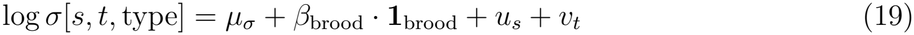

The prior on *µ*_*σ*_ is centered on log(0.08), corresponding to a prior median detection scale of 80 m, consistent with published half-normal estimates for grouse surveyed by foot transect. The standard deviation of 1 on the log scale places 95 % prior probability on *σ* ∈ (11, 570) m, wide enough to accommodate substantial site-year variation while excluding implausible values.

Detection probability varied substantially across sites and years (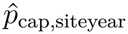 range: 0.30– 0.62), with brood groups detected at slightly longer distances than non-brood groups (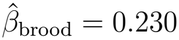, 95 % CrI 0.122–0.337), justifying the brood-composition-weighted detection probability used in density estimation. Posterior predictive checks confirmed adequate model fit to encounter frequencies and young counts by adult category; the distance-bin statistic indicated clustering of encounters within transects (Appendix A6).

#### A2.3 Distance bin probabilities and encounter likelihood

The probability that a detected encounter falls in distance bin *k* is:

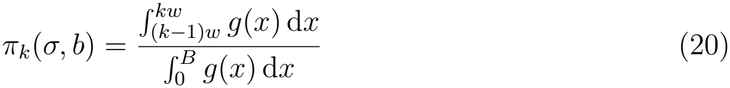

computed numerically within JAGS for each site-year-type combination (8 bins, truncation at *B* = 0.2 km). The distance bin of encounter *i* follows a categorical distribution over (*π*_1_, …, *π*_8_). The site-year average detectability is a brood-composition-weighted average of brood and non-brood detection probabilities:

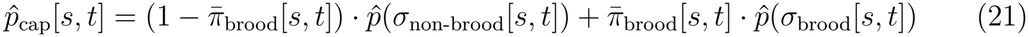

where 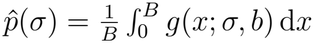,

#### A2.4 Adult group size

Category probabilities are modeled via a multinomial logit (softmax), with category 1 (single adult) as the reference:

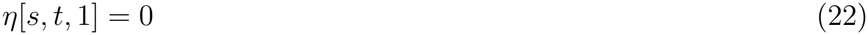

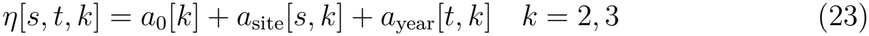

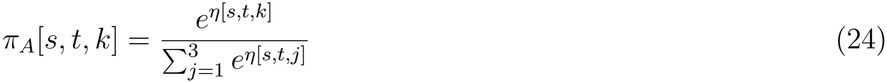

The expected number of adults per group is *λ*_*A*_[*s, t*] = 1 · *π*_*A*_[*s, t*, 1] + 2 · *π*_*A*_[*s, t*, 2] + *k*_3_ · *π*_*A*_[*s, t*, 3], where *k*_3_ = 3.44 is the fixed mean size of groups in the 3+ category. Adult density is *D*_*A*_[*s, t*] = *D*_grp_[*s, t*] · *λ*_*A*_[*s, t*].

#### A2.5 Young counts: two-part model and zeros trick

The number of young per group is modeled in two parts: a logistic model for brood occurrence (*Y >* 0) and a zero-truncated negative binomial for positive counts *Y* | *Y >* 0. Both components have adult-category-specific intercepts, with site and year random effects (*σ*_*πY,s*_, *σ*_*πY,t*_, *σ*_*Y* +,*s*_, *σ*_*Y* +,*t*_; Table 8):

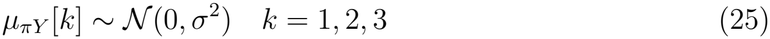

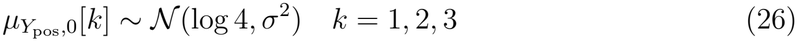

Because JAGS lacks a built-in zero-truncated negative binomial, we used the zeros trick to evaluate the correct log-likelihood contribution. For zero-young encounters, the log-likelihood is log(1 − *π*_brood_[*s, t, k*]). For positive encounters it is: log(*π*_brood_) + log *p*_NB_(*Y*_*i*_) − log(1−*p*_NB_(0)), where the truncation correction − log(1−*p*_NB_(0)) renormalizes the negative binomial to exclude zero. Omitting this correction yields biased estimates of the positive brood size distribution.

Young per adult is the product of two distinct processes, the proportion of females that successfully raise a brood and the size of the broods they raise, which need not move together. In this study, we modeled them separately, with a logistic component for brood occurrence and a zero-truncated negative binomial for positive brood size, rather than collapsing both into a single index. This two-part structure recovered a clear effect of productivity on population change in the Gompertz model. Because productivity is the principal driver of realized dynamics in this system, a less precise productivity index can obscure the signal that matters most for both inference and management. Young per adult has been estimated differently in different studies (Steen, Steen, et al. 1988; Hörnell-Willebrand, Marcström, et al. 2006; Breisjøberget et al. 2018).

#### A2.6 Propagating detection uncertainty to dynamic models

The static posterior for each site-year is summarized on the logit scale:

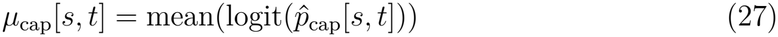

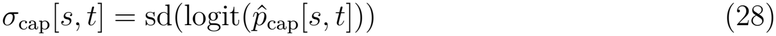

These are saved to mu_cap.csv and sd_cap.csv and used as an informative prior in the dynamic models:

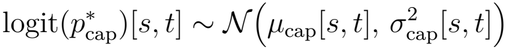

This propagates detection uncertainty without refitting the full observation sub-model; the encounter counts in the dynamic models can still update 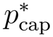. The logit-scale posteriors were verified as approximately normal by visual inspection; the observed range 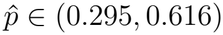 lies well within the linear region of the logit transform, supporting the Gaussian approximation.

### A3. Hazard model details

#### A3.1 Natural hazard prior

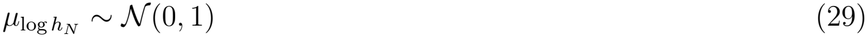

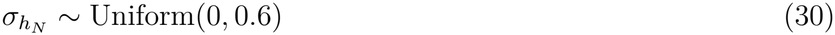

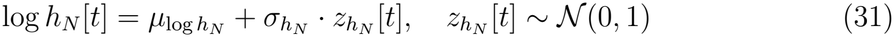

The log-normal structure ensures strict positivity and multiplicative year deviations on the hazard scale.

#### A3.2 Harvest hazard and process priors

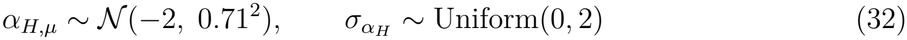

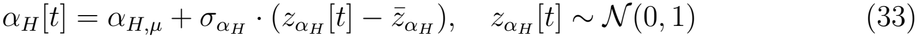

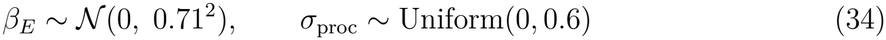

#### A3.3 Bag likelihood

Expected bag size at open sites is:

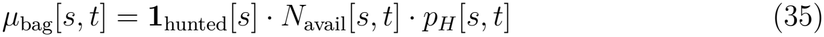

where *N*_avail_[*s, t*] = Area[*s*] · (*D*_*A*_[*s, t*] + *κ*_*Y*_ · *D*_*Y*_ [*s, t*]) adjusts for differential availability of juveniles relative to adults via the parameter *κ*_*Y*_, and *p*_*H*_[*s, t*] = 1 − exp(−*h*_*H*_[*s, t*]) is the implied harvest probability. Bag counts follow a negative binomial distribution with mean *µ*_bag_ and overdispersion parameter *r*_*C*_ ∼ Uniform(0.5, 60); the availability ratio has prior log *κ*_*Y*_ ∼ *N* (0, 0.5^2^).

#### A3.4 Derived management scalars

Year-averaged survival summaries are computed by averaging over year-specific quantities rather than using a point estimate. The absolute difference in total survival between effort one SD above the center of the standardized scale and effort at the center is:

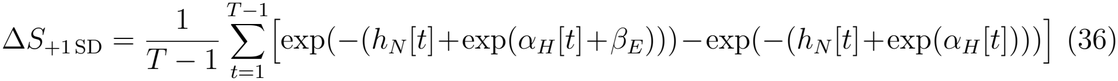

Table 2 instead evaluates survival at the mean hazards (Methods).

### A4. Gompertz model details

#### A4.0 What the Gompertz model can and cannot resolve regarding effort

The Gompertz model reproduced the observed dynamics well (paragraph 2.3), but this does not imply that its effort and feedback parameters are identified. A model with site intercepts, a productivity covariate and a process variance partitions the variance among *α*[*s*], *ϕ* and *β*_*E*_. First, *β*_*E*_ is identified only from year-to-year variation in effort within the three open sites; it measures the net one-year response of growth, including any compensation, but not a sustained difference in density level, which is absorbed by *α*[*s*] and confounded with site differences. Second, *ϕ* is weakly identified from eleven transitions per site; we assessed this by comparing its posterior with its prior (A4.2). Third, the first-order autoregression does not include delayed feedback, such as lagged numerical responses of predators. We therefore report the Gompertz estimates as a description of realized short-term dynamics, not as a basis for effort thresholds. Models that separate the sources of negative density feedback using these data are developed elsewhere (in prep.).

#### A4.1 Site intercept hierarchy

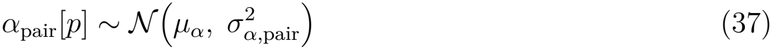

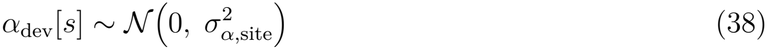

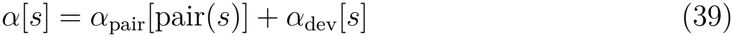

This pair-level structure respects the hierarchical design and allows partial pooling within pairs while capturing genuine site-level variation.

At mean productivity, equilibrium adult densities ranged from 4.3 to 8.4 adults km^−2^ among sites (open relative to closed: A +18 %, B +87 %, C −44 %; posterior summaries in Table 8). The estimated density dependence (*ϕ* = 0.321) implies that about two-thirds (1 − *ϕ* = 0.68) of a deviation from the site’s equilibrium level is removed per year.

#### A4.2 Prior on the autoregressive coefficient

The autoregressive coefficient *ϕ* is constrained to (0, 1) via a logistic transformation of a normally distributed raw parameter:

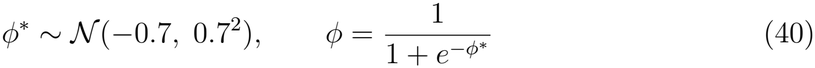

This prior places 95 % of its probability in the range *ϕ* ∈ (0.11, 0.66), consistent with strong mean-reverting dynamics, while excluding the boundary values 0 (complete return to the site’s equilibrium level within one year) and 1 (random walk, no density dependence). The posterior (median 0.32, 95 % CrI 0.13–0.55) is only modestly narrower than the prior (median 0.33, 95 % interval 0.11–0.66), so the data provide limited information on *ϕ* beyond the prior.

#### A4.3 Productivity covariate

The productivity index is 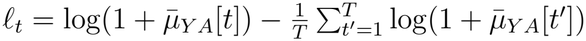, where 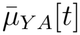 [*t*] is the number of young per adult averaged across sites in year *t*. It is centered over years within each MCMC sample to ensure orthogonality with the intercept.

#### A4.4 Long-run effect of constant effort

Under constant effort, the stationary mean of log adult density on an open site shifts by *β*_*E*_*E*_*sc*_*/*(1 − *ϕ*). Computed per posterior draw, the long-run change in median density at 3 hd km^−2^ relative to no harvest is −13 % (95 % CrI −54 to +42 %). Zero effort lies below the lowest observed effort (0.16 hd km^−2^), so this contrast extrapolates the effort slope.

### A5. Effort standardization

Predictions at the management thresholds used the same standardized scale as model fitting. Raw effort *E* (hunter-days per 10 km^2^) was transformed as:

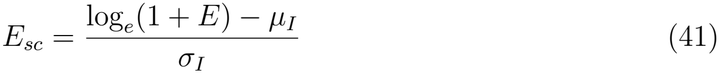

where *µ*_*I*_ = 2.594 and *σ*_*I*_ = 0.750 are the mean and standard deviation of log_*e*_(1 + *E*) across all open site-years in the observed data. The management reference thresholds of 1 and 3 hd km^−2^ correspond to 10 and 30 hd per 10 km^2^, respectively, yielding:

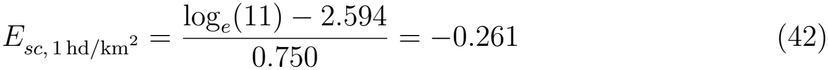

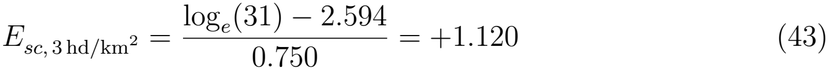

Both values fall within the observed range of standardized effort (−2.16 to +2.14), confirming that predictions for the management reference levels of 1 and 3 hd km^−2^ are interpolations rather than extrapolations.

### A6. Posterior predictive checks

We checked each submodel of the static model with posterior predictive statistics (Table 5): a Freeman–Tukey discrepancy for the encounter counts per site-year, *χ*^2^ discrepancies for the adult group-size categories (*T*_*a*_) and for the distance-bin counts (*T*_*d*_), and summaries of the young counts by number of adults. The encounter-rate and group-size checks were well calibrated (*p* = 0.51 and 0.50). The distance-bin discrepancy *T*_*d*_ showed a large observed value relative to replicated values (*p <* 0.0001). We attribute this to spatial clustering of encounters within transects rather than to bias in the detection function. Clustering of encounters inflates the variance of bin counts rather than shifting their expected distribution, so we expect little bias in 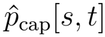, although its uncertainty is probably understated. We therefore accepted the model for inference.

The young-count checks were well calibrated (*p* = 0.29–0.52; Table 5), except that the proportion of encounters with eight or more young was under-predicted for groups with one adult (*p* = 0.012) and slightly for groups with two adults (*p* = 0.100), suggesting that the largest broods are not fully captured by the young-count model.

For the Hazard dynamic model, we evaluated whether the process variance *σ*_proc_ was correctly specified and whether the bag likelihood adequately characterizes the distribution of bag counts. The process-level checks use a propagated replicated trajectory initialized at the latent states. At each subsequent step, the replicated trajectory uses its own previous state but the *fitted* survival *S*[*s, t*]. By using the fitted *S*[*s, t*] for both observed and replicated trajectories, the PPC isolates *the adequacy of the process noise specification* from any potential misspecification in the survival model itself.

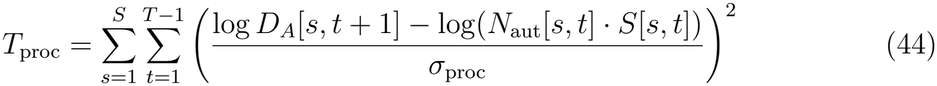

The observed bags were compared with bags replicated from the model using the Freeman–Tukey discrepancy between bag and expected bag. The discrepancy was smaller for the observed than for the replicated bags (posterior means 51.0 and 55.0; *p* = 0.56), indicating adequate fit, with observed bags slightly less dispersed around their expectation than the negative binomial bag likelihood allows.

We evaluated whether the model reproduces the observed difference in mean log-growth between high- and low-effort open-site years. The model-generated contrast is slightly smaller in magnitude than the observed contrast (Δ*r*_rep_ = −0.142 versus Δ*r*_obs_ = −0.158; Table 6), indicating that a small part of the observed growth-rate difference between high- and low-effort site-years is not fully captured by the Hazard model’s parameterization of the effort effect. This is consistent with the one-step residuals in Appendix A7.

In the Gompertz dynamic model, the process-variance check was:

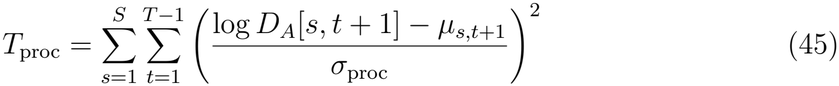

For both dynamic models, the process variance PPC confirmed adequate calibration (*p* ≈ 0.53). However, adequate calibration does not imply that individual parameters are well identified (Appendix A4).

The process standard deviation *σ*_proc_ differs substantially between the models: 0.199 (95 % CrI 0.142–0.274) in the Hazard model versus 0.113 (95 % CrI 0.022–0.193) in the Gompertz model. This reflects the difference in how the two models handle annual variation. The Gompertz model explicitly accounts for density dependence and productivity as drivers of annual growth, absorbing a substantial fraction of annual variation into its deterministic part.

The Hazard model operates on a mortality scale and does not directly include density dependence, so a larger part of the unexplained variation is assigned to the process error term. The larger *σ*_proc_ in the Hazard model also explains why the replicated data showed greater variation in growth rates than observed (mean absolute and mean squared log-growth, 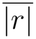 and 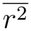, Table 6).

The posterior means of young per adult (*µ*_*Y A*_) from the Hazard and Gompertz models were highly correlated across site-years (*r* = 0.99), and the yearly productivity signal agreed closely between the two models (Figure 5). The shared young submodel is therefore robust to the choice of population process; this check does not address whether the two models agree on survival or on the effort effect.

**Figure 5:**
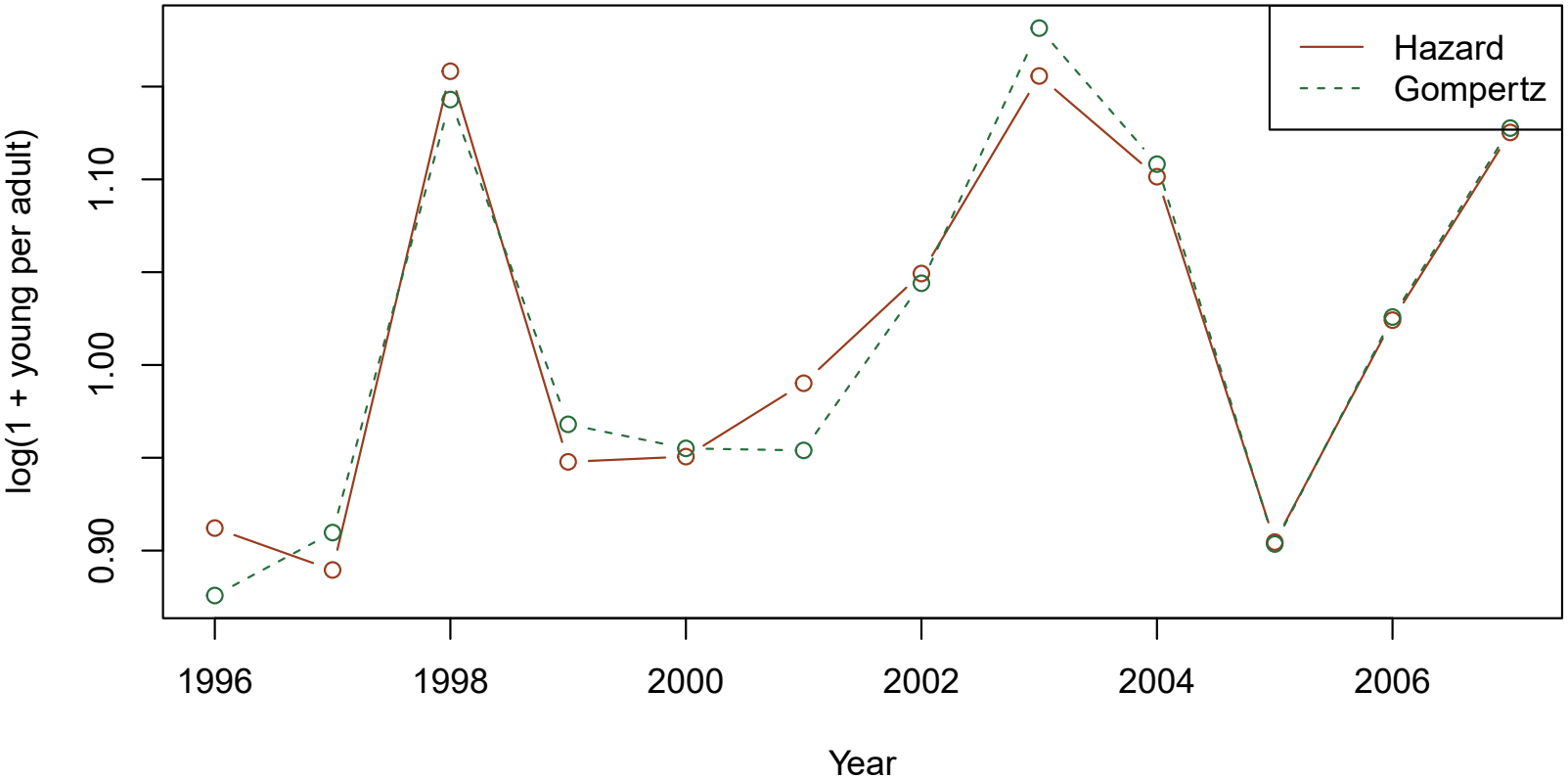
Yearly productivity signal estimated by the Hazard (solid red) and Gompertz (dashed green) models: posterior mean of 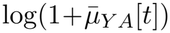, where 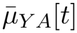 is the number of young per adult averaged over sites (the uncentered productivity index; *ℓ*_*t*_ in Eq. 6 is this quantity centered over years). Both models share the same young submodel, and the close agreement (*r* = 0.99) shows that productivity estimates are nearly independent of the choice of population process model.

### A7. Sensitivity check: shared natural-hazard assumption

#### Rationale

The process equation of the Hazard model imposes a single year-specific natural hazard *h*_*N*_ [*t*] on all sites. However, compensatory predation or spatially structured predator territories could in principle generate higher natural mortality in open sites, which would cause the model to attribute genuine natural mortality to harvest hazard, inflating *β*_*E*_. We used one-step-ahead residuals to test this post hoc.

For each posterior sample *i* and each site-year transition (*s, t* → *t* + 1), the standardized residual is

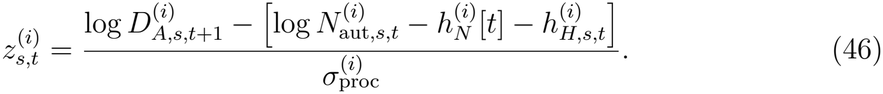

Posterior mean residuals 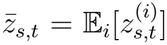 were summarized by open/closed group and by open-site pairing.

One-step-ahead residuals from the Hazard model tended to be positive in open sites relative to closed sites (mean difference = +0.17 z-units, *P* (open *>* closed) = 0.86, 95 % CrI −0.13 to +0.49), positive in pairs A and B and negative in pair C. Positive residuals on open sites argue against the concern that higher natural mortality there inflates *β*_*E*_; they correspond to the gain estimated in paragraph 2.3. The Pearson correlation between *E*_*sc*_ and 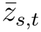 in open sites was *r* = −0.32 (*p* = 0.065, *n* = 5,000 permutations), which suggests, though not significantly, that high-effort years tend to over-predict next-year density, so that *h*_*H*_ may under-capture the effort signal.

The pattern is weak and cannot be separated from effort-related natural mortality without independent survival data.

### A8. Full posterior parameter summaries

