## Supplementary Material: Hunter effort increases willow ptarmigan mortality, but the cost is not visible in abundance: evaluating effort thresholds for for "Hunter effort increases willow ptarmigan mortality, but the cost is not visible in abundance: evaluating effort thresholds for harvest management"

### Supplementary Material: Evaluating effort as a management tool for small game. Using distance sampling with two dynamic models, the Gompertz and Hazard.

Tomas

May 29, 2026

Development notes (all three) ✓JAGS model code ✓R analysis scripts ✓Full convergence diagnostics ✓Posterior predictive checks ✓Tables of key parameters and derived quantities ✓Figures of model fit and key results.

#### 1 Development notes from project Workbook

See separate documents: Gommpertz_details.pdf, Hazard_details.pdf, and Static_details.pdf for detailed notes on model development, MCMC settings, convergence diagnostics, and posterior predictive checks for each of the three models.

#### 2 Jags code for Static, Hazard adn Gompertz models

#### 3 R analysis scripts

#### 4 Full convergence diagnostics

Hazard model:
All Rhat < 1.05 – good convergence
n.eff < 500:
param Rhat n.eff
alpha_H_mu 1.014362 476

Gompertz model:
All Rhat < 1.05 – good convergence
All n.eff >= 500

**Table 1:** Full posterior summary for the hazard state-space model. Mean, SD, 2.5%, median, and 97.5% quantiles of the marginal posterior distribution. $\hat{R}$: Gelman–Rubin convergence statistic (values $< 1.1$ indicate convergence). $n_{\text{eff}}$: effective sample size. Overlap 0: whether the 95% credible interval includes zero. $f$: posterior probability that the parameter has the same sign as the posterior mean.

| Parameter | Mean | SD | 2.5% | Median | 97.5% | $\hat{R}$ | $n_{\text{eff}}$ | Overlap 0 | $f$ |
|---|---|---|---|---|---|---|---|---|---|
| *Process parameters* | | | | | | | | | |
| $\beta_E$ (effort effect on harvest hazard) | 0.826 | 0.081 | 0.665 | 0.826 | 0.983 | 1.001 | 11,263 | no | 1.000 |
| $\sigma_{\text{proc}}$ (process SD, log scale) | 0.196 | 0.033 | 0.137 | 0.194 | 0.265 | 1.001 | 13,802 | no | 1.000 |
| *Natural mortality hazard* | | | | | | | | | |
| $\mu_{\log h_N}$ (grand mean log natural hazard) | -0.176 | 0.064 | -0.314 | -0.173 | -0.057 | 1.002 | 3,403 | no | 0.996 |
| $\sigma_{h_N}$ (year-level SD, log scale) | 0.148 | 0.083 | 0.013 | 0.140 | 0.339 | 1.001 | 9,912 | no | 1.000 |
| *Harvest hazard* | | | | | | | | | |
| $\alpha_{H,\mu}$ (harvest baseline mean) | -2.369 | 0.289 | -2.954 | -2.362 | -1.822 | 1.014 | 476 | no | 1.000 |
| $\sigma_{\alpha_H}$ (year-level SD) | 0.667 | 0.181 | 0.396 | 0.638 | 1.097 | 1.004 | 1,871 | no | 1.000 |
| *Year-level harvest hazard baseline* $\alpha_H[t]$ | | | | | | | | | |
| Year 1 (1996) | -1.772 | 0.274 | -2.345 | -1.761 | -1.265 | 1.010 | 701 | no | 1.000 |
| Year 2 (1997) | -2.214 | 0.297 | -2.824 | -2.206 | -1.658 | 1.009 | 783 | no | 1.000 |
| Year 3 (1998) | -2.142 | 0.305 | -2.771 | -2.132 | -1.580 | 1.009 | 729 | no | 1.000 |
| Year 4 (1999) | -2.250 | 0.256 | -2.788 | -2.238 | -1.783 | 1.010 | 682 | no | 1.000 |
| Year 5 (2000) | -2.067 | 0.259 | -2.609 | -2.054 | -1.597 | 1.010 | 684 | no | 1.000 |
| Year 6 (2001) | -1.820 | 0.264 | -2.376 | -1.807 | -1.340 | 1.011 | 621 | no | 1.000 |
| Year 7 (2002) | -2.155 | 0.271 | -2.723 | -2.142 | -1.660 | 1.011 | 598 | no | 1.000 |
| Year 8 (2003) | -2.438 | 0.277 | -3.014 | -2.425 | -1.933 | 1.012 | 588 | no | 1.000 |
| Year 9 (2004) | -3.694 | 0.288 | -4.287 | -3.684 | -3.157 | 1.010 | 694 | no | 1.000 |
| Year 10 (2005) | -3.306 | 0.265 | -3.856 | -3.297 | -2.811 | 1.009 | 749 | no | 1.000 |
| Year 11 (2006) | -2.358 | 0.268 | -2.919 | -2.347 | -1.867 | 1.010 | 648 | no | 1.000 |

*continued on next page*


3

*continued from previous page*

| Parameter | Mean | SD | 2.5% | Median | 97.5% | $\hat{R}$ | $n_{\mathrm{eff}}$ | Overlap 0 | $f$ |
|---|---|---|---|---|---|---|---|---|---|
| Year 12 (2007) | -2.673 | 0.289 | -3.268 | -2.662 | -2.138 | 1.010 | 676 | no | 1.000 |
| *Bag likelihood* | | | | | | | | | |
| $\kappa_Y$ (juvenile availability ratio) | 1.186 | 0.434 | 0.570 | 1.108 | 2.252 | 1.013 | 538 | no | 1.000 |
| $\log \kappa_Y$ (log scale) | 0.109 | 0.352 | -0.563 | 0.103 | 0.812 | 1.014 | 495 | yes | 0.617 |
| $r_C$ (bag NB overdispersion) | 47.003 | 9.713 | 24.589 | 48.982 | 59.528 | 1.000 | 59,694 | no | 1.000 |
| *Young sub-model* | | | | | | | | | |
| $\mu_{\pi Y}[1]$ (brood occurrence intercept, 1 adult) | -0.203 | 0.156 | -0.517 | -0.203 | 0.109 | 1.002 | 3,793 | yes | 0.910 |
| $\mu_{\pi Y}[2]$ (brood occurrence intercept, 2 adults) | 0.440 | 0.155 | 0.129 | 0.440 | 0.749 | 1.002 | 3,751 | no | 0.995 |
| $\mu_{\pi Y}[3]$ (brood occurrence intercept, 3+ adults) | -0.648 | 0.205 | -1.058 | -0.647 | -0.244 | 1.001 | 6,272 | no | 0.999 |
| $\mu_{Y+,0}[1]$ (positive brood size intercept, 1 adult) | 1.492 | 0.076 | 1.335 | 1.494 | 1.638 | 1.004 | 2,163 | no | 1.000 |
| $\mu_{Y+,0}[2]$ (positive brood size intercept, 2 adults) | 1.604 | 0.074 | 1.450 | 1.606 | 1.746 | 1.005 | 1,954 | no | 1.000 |
| $\mu_{Y+,0}[3]$ (positive brood size intercept, 3+ adults) | 1.937 | 0.098 | 1.739 | 1.938 | 2.125 | 1.003 | 3,191 | no | 1.000 |
| $\log r_Y$ (young count NB dispersion, log scale) | 1.582 | 0.078 | 1.432 | 1.581 | 1.736 | 1.000 | 49,734 | no | 1.000 |
| $\sigma_{\pi Y,t}$ (brood occurrence year SD) | 0.332 | 0.094 | 0.194 | 0.316 | 0.559 | 1.000 | 28,667 | no | 1.000 |
| $\sigma_{\pi Y,s}$ (brood occurrence site SD) | 0.352 | 0.105 | 0.183 | 0.339 | 0.571 | 1.000 | 22,654 | no | 1.000 |
| $\sigma_{Y+,t}$ (brood size year SD) | 0.108 | 0.034 | 0.057 | 0.103 | 0.191 | 1.000 | 23,882 | no | 1.000 |
| $\sigma_{Y+,s}$ (brood size site SD) | 0.141 | 0.063 | 0.060 | 0.126 | 0.309 | 1.002 | 3,977 | no | 1.000 |
| *Adult group size sub-model* | | | | | | | | | |
| $a_0[2]$ (log-odds of 2 adults vs 1, grand mean) | 0.187 | 0.200 | -0.233 | 0.193 | 0.569 | 1.009 | 1,352 | yes | 0.866 |
| $a_0[3]$ (log-odds of 3+ adults vs 1, grand mean) | -1.914 | 0.360 | -2.520 | -1.958 | -1.045 | 1.004 | 2,211 | no | 1.000 |
| *Annual productivity index* $\log \bar{\mu}_{YA}[t]$ | | | | | | | | | |
| Year 1 (1996) | 0.819 | 0.051 | 0.720 | 0.819 | 0.918 | 1.000 | 55,180 | no | 1.000 |
| Year 2 (1997) | 0.781 | 0.059 | 0.664 | 0.782 | 0.897 | 1.000 | 64,933 | no | 1.000 |
| Year 3 (1998) | 1.092 | 0.040 | 1.013 | 1.092 | 1.172 | 1.000 | 57,669 | no | 1.000 |
| Year 4 (1999) | 0.877 | 0.037 | 0.805 | 0.877 | 0.950 | 1.000 | 60,767 | no | 1.000 |
| Year 5 (2000) | 0.855 | 0.038 | 0.781 | 0.855 | 0.929 | 1.000 | 65,598 | no | 1.000 |
| Year 6 (2001) | 0.931 | 0.039 | 0.855 | 0.931 | 1.008 | 1.000 | 132,464 | no | 1.000 |
| Year 7 (2002) | 0.973 | 0.039 | 0.895 | 0.972 | 1.050 | 1.000 | 61,770 | no | 1.000 |

*continued on next page*


4

*continued from previous page*

| Parameter | Mean | SD | 2.5% | Median | 97.5% | $\hat{R}$ | $n_{\text{eff}}$ | Overlap 0 | $f$ |
|---|---|---|---|---|---|---|---|---|---|
| Year 8 (2003) | 1.096 | 0.038 | 1.023 | 1.096 | 1.171 | 1.000 | 54,619 | no | 1.000 |
| Year 9 (2004) | 1.047 | 0.035 | 0.978 | 1.047 | 1.117 | 1.000 | 37,112 | no | 1.000 |
| Year 10 (2005) | 0.822 | 0.038 | 0.747 | 0.822 | 0.897 | 1.000 | 49,207 | no | 1.000 |
| Year 11 (2006) | 0.920 | 0.045 | 0.832 | 0.920 | 1.008 | 1.000 | 166,099 | no | 1.000 |
| Year 12 (2007) | 1.026 | 0.045 | 0.940 | 1.026 | 1.114 | 1.000 | 117,023 | no | 1.000 |
| *Derived management quantities* | | | | | | | | | |
| $\text{relSH}_{+1\,\text{SD}}$ (relative harvest survival at +1 SD effort) | 0.882 | 0.038 | 0.795 | 0.887 | 0.942 | 1.012 | 626 | no | 1.000 |
| $\text{relSH}_{-1\,\text{SD}}$ (relative harvest survival at −1 SD effort) | 1.057 | 0.018 | 1.029 | 1.054 | 1.098 | 1.014 | 524 | no | 1.000 |
| $\Delta S_{+1\,\text{SD}}$ (absolute total survival change at +1 SD effort) | -0.046 | 0.014 | -0.077 | -0.044 | -0.023 | 1.011 | 633 | no | 1.000 |
| *Posterior predictive check statistics* | | | | | | | | | |
| $T_{\text{proc,obs}}$ (process $\chi^2$, observed) | 65.000 | 11.381 | 44.643 | 64.307 | 89.080 | 1.000 | 118,673 | no | 1.000 |
| $T_{\text{proc,rep}}$ (process $\chi^2$, replicated) | 65.952 | 11.507 | 45.372 | 65.294 | 90.312 | 1.000 | 90,016 | no | 1.000 |
| $\overline{\lvert r \rvert}_{\text{obs}}$ (mean absolute log-growth, observed) | 0.181 | 0.020 | 0.144 | 0.180 | 0.222 | 1.000 | 12,645 | no | 1.000 |
| $\overline{\lvert r \rvert}_{\text{rep}}$ (mean absolute log-growth, replicated) | 0.216 | 0.030 | 0.162 | 0.214 | 0.281 | 1.000 | 12,906 | no | 1.000 |
| $\overline{r^2}_{\text{obs}}$ (mean squared log-growth, observed) | 0.049 | 0.010 | 0.032 | 0.048 | 0.071 | 1.001 | 12,591 | no | 1.000 |
| $\overline{r^2}_{\text{rep}}$ (mean squared log-growth, replicated) | 0.073 | 0.020 | 0.041 | 0.071 | 0.120 | 1.000 | 13,297 | no | 1.000 |
| $\bar{r}_{\text{hi,obs}}$ (mean log-growth, high effort years, observed) | -0.051 | 0.022 | -0.094 | -0.050 | -0.007 | 1.000 | 16,214 | no | 0.989 |
| $\bar{r}_{\text{lo,obs}}$ (mean log-growth, low effort years, observed) | 0.067 | 0.019 | 0.030 | 0.067 | 0.104 | 1.000 | 21,932 | no | 1.000 |
| $\bar{r}_{\text{hi,rep}}$ (mean log-growth, high effort years, replicated) | -0.046 | 0.067 | -0.177 | -0.047 | 0.088 | 1.001 | 7,319 | yes | 0.765 |
| $\bar{r}_{\text{lo,rep}}$ (mean log-growth, low effort years, replicated) | 0.020 | 0.057 | -0.094 | 0.021 | 0.131 | 1.000 | 75,614 | yes | 0.646 |
| $\Delta r_{\text{obs}}$ (hi−lo effort contrast, observed) | -0.117 | 0.035 | -0.186 | -0.117 | -0.050 | 1.000 | 16,457 | no | 1.000 |
| $\Delta r_{\text{rep}}$ (hi−lo effort contrast, replicated) | -0.067 | 0.083 | -0.227 | -0.068 | 0.100 | 1.000 | 12,917 | yes | 0.796 |
| $T_{\text{bag,obs}}$ (bag Freeman-Tukey, observed) | 37.884 | 13.438 | 19.353 | 35.428 | 70.755 | 1.000 | 48,988 | no | 1.000 |
| $T_{\text{bag,rep}}$ (bag Freeman-Tukey, replicated) | 45.965 | 17.336 | 22.723 | 42.715 | 88.619 | 1.000 | 80,808 | no | 1.000 |
| $p_{\text{bag}}$ (bag PPC Bayesian $p$-value) | 0.685 | 0.465 | 0.000 | 1.000 | 1.000 | 1.000 | 143,089 | yes | 1.000 |


5

**Table 2:** Full posterior summary for the Gompertz state-space model. See Table 1 caption for column definitions.

| Parameter | Mean | SD | 2.5% | Median | 97.5% | $\hat{R}$ | $n_{\text{eff}}$ | Overlap 0 | $f$ |
|---|---|---|---|---|---|---|---|---|---|
| *Structural parameters* | | | | | | | | | |
| $\beta_E$ (effort effect on log growth rate) | -0.012 | 0.036 | -0.081 | -0.013 | 0.059 | 1.001 | 5,889 | yes | 0.640 |
| $\phi$ (density feedback coefficient) | 0.327 | 0.110 | 0.131 | 0.323 | 0.553 | 1.002 | 2,925 | no | 1.000 |
| $\gamma$ (productivity effect) | 0.934 | 0.240 | 0.481 | 0.928 | 1.430 | 1.001 | 4,591 | no | 1.000 |
| $\sigma_{\text{proc}}$ (process SD) | 0.096 | 0.040 | 0.015 | 0.097 | 0.173 | 1.013 | 601 | no | 1.000 |
| *Site equilibrium densities* | | | | | | | | | |
| $\mu_\alpha$ (grand mean log-density) | 1.899 | 0.131 | 1.608 | 1.906 | 2.147 | 1.000 | 40,249 | no | 1.000 |
| $\sigma_{\alpha,\text{pair}}$ (pair-level SD) | 0.126 | 0.116 | 0.004 | 0.093 | 0.442 | 1.002 | 5,086 | no | 1.000 |
| $\sigma_{\alpha,\text{site}}$ (site-level SD) | 0.185 | 0.081 | 0.076 | 0.168 | 0.387 | 1.005 | 1,742 | no | 1.000 |
| *Site-specific equilibria* $\alpha[s]$ | | | | | | | | | |
| $\alpha[1]$ (site A0) | 2.052 | 0.046 | 1.960 | 2.052 | 2.142 | 1.000 | 17,728 | no | 1.000 |
| $\alpha[2]$ (site A1) | 1.844 | 0.052 | 1.737 | 1.846 | 1.944 | 1.001 | 15,092 | no | 1.000 |
| $\alpha[3]$ (site B0) | 1.795 | 0.061 | 1.672 | 1.797 | 1.908 | 1.002 | 3,063 | no | 1.000 |
| $\alpha[4]$ (site B1) | 2.103 | 0.050 | 2.005 | 2.103 | 2.202 | 1.001 | 6,857 | no | 1.000 |
| $\alpha[5]$ (site C0) | 1.792 | 0.059 | 1.672 | 1.793 | 1.901 | 1.001 | 5,199 | no | 1.000 |
| $\alpha[6]$ (site C1) | 1.898 | 0.052 | 1.789 | 1.900 | 1.994 | 1.001 | 5,729 | no | 1.000 |
| *Annual productivity index* $\log \bar{\mu}_{YA}[t]$ | | | | | | | | | |
| Year 1 (1996) | 0.791 | 0.049 | 0.695 | 0.791 | 0.887 | 1.001 | 14,128 | no | 1.000 |
| Year 2 (1997) | 0.784 | 0.056 | 0.671 | 0.785 | 0.891 | 1.000 | 13,342 | no | 1.000 |
| Year 3 (1998) | 1.078 | 0.039 | 1.002 | 1.077 | 1.155 | 1.001 | 12,581 | no | 1.000 |
| Year 4 (1999) | 0.892 | 0.036 | 0.821 | 0.893 | 0.961 | 1.000 | 20,169 | no | 1.000 |
| Year 5 (2000) | 0.851 | 0.036 | 0.780 | 0.851 | 0.920 | 1.000 | 19,997 | no | 1.000 |
| Year 6 (2001) | 0.898 | 0.038 | 0.825 | 0.897 | 0.972 | 1.001 | 9,504 | no | 1.000 |
| Year 7 (2002) | 0.961 | 0.037 | 0.889 | 0.961 | 1.034 | 1.000 | 22,028 | no | 1.000 |
| Year 8 (2003) | 1.114 | 0.036 | 1.044 | 1.114 | 1.185 | 1.000 | 28,105 | no | 1.000 |
| Year 9 (2004) | 1.060 | 0.034 | 0.994 | 1.060 | 1.127 | 1.001 | 6,277 | no | 1.000 |

*continued on next page*



9

*continued from previous page*

| Parameter | Mean | SD | 2.5% | Median | 97.5% | $\hat{R}$ | $n_{\text{eff}}$ | Overlap 0 | $f$ |
|---|---|---|---|---|---|---|---|---|---|
| Year 10 (2005) | 0.828 | 0.037 | 0.756 | 0.828 | 0.901 | 1.000 | 23,619 | no | 1.000 |
| Year 11 (2006) | 0.928 | 0.042 | 0.845 | 0.928 | 1.011 | 1.000 | 31,121 | no | 1.000 |
| Year 12 (2007) | 1.028 | 0.045 | 0.941 | 1.028 | 1.117 | 1.000 | 67,088 | no | 1.000 |
| *PPC statistics* | | | | | | | | | |
| $T_{\text{proc,obs}}$ (observed process chi-sq) | 64.978 | 11.403 | 44.579 | 64.321 | 89.213 | 1.000 | 82,671 | no | 1.000 |
| $T_{\text{proc,rep}}$ (replicated process chi-sq) | 65.996 | 11.497 | 45.474 | 65.312 | 90.399 | 1.000 | 168,000 | no | 1.000 |
| $\overline{\lvert r \rvert}_{\text{obs}}$ (mean abs growth rate, obs) | 0.152 | 0.026 | 0.103 | 0.151 | 0.205 | 1.005 | 1,390 | no | 1.000 |
| $\overline{\lvert r \rvert}_{\text{rep}}$ (mean abs growth rate, rep) | 0.151 | 0.030 | 0.100 | 0.149 | 0.217 | 1.004 | 1,863 | no | 1.000 |
| $\Delta r_{\text{obs}}$ (hi−lo effort growth contrast, obs) | -0.113 | 0.034 | -0.183 | -0.112 | -0.050 | 1.001 | 6,136 | no | 1.000 |
| $\Delta r_{\text{rep}}$ (hi−lo effort growth contrast, rep) | -0.097 | 0.043 | -0.183 | -0.097 | -0.010 | 1.000 | 23,588 | no | 0.984 |


7

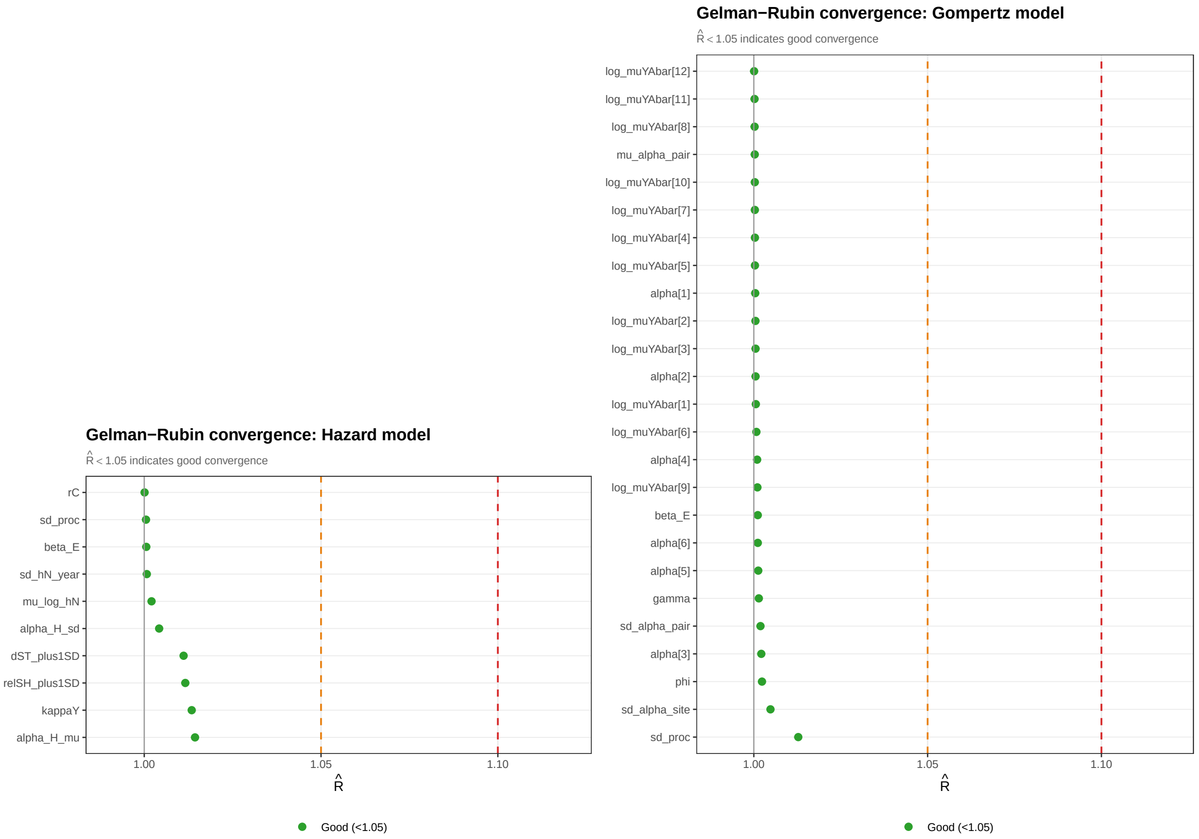


**(a)** Hazard Model

**(b)** Gompertz Model

**Figure 1:** Graphs showing Rhat values for all parameters in the Hazard and Gompertz models. All values are below 1.05, indicating good convergence.

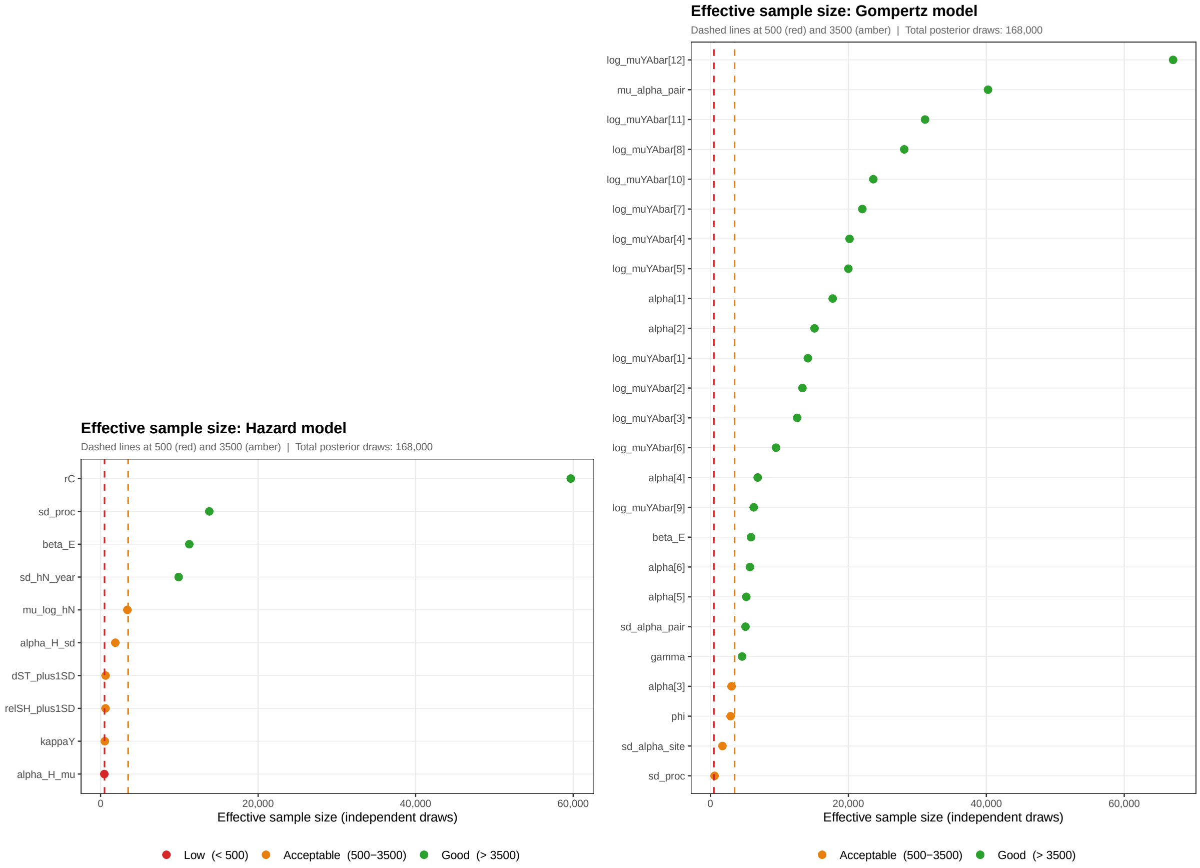


**(a)** Hazard Model　　**(b)** Gompertz Model

**Figure 2:** Graphs showing n.eff values for all parameters in the Hazard and Gompertz models. All except one value is above 500, indicating good convergence.

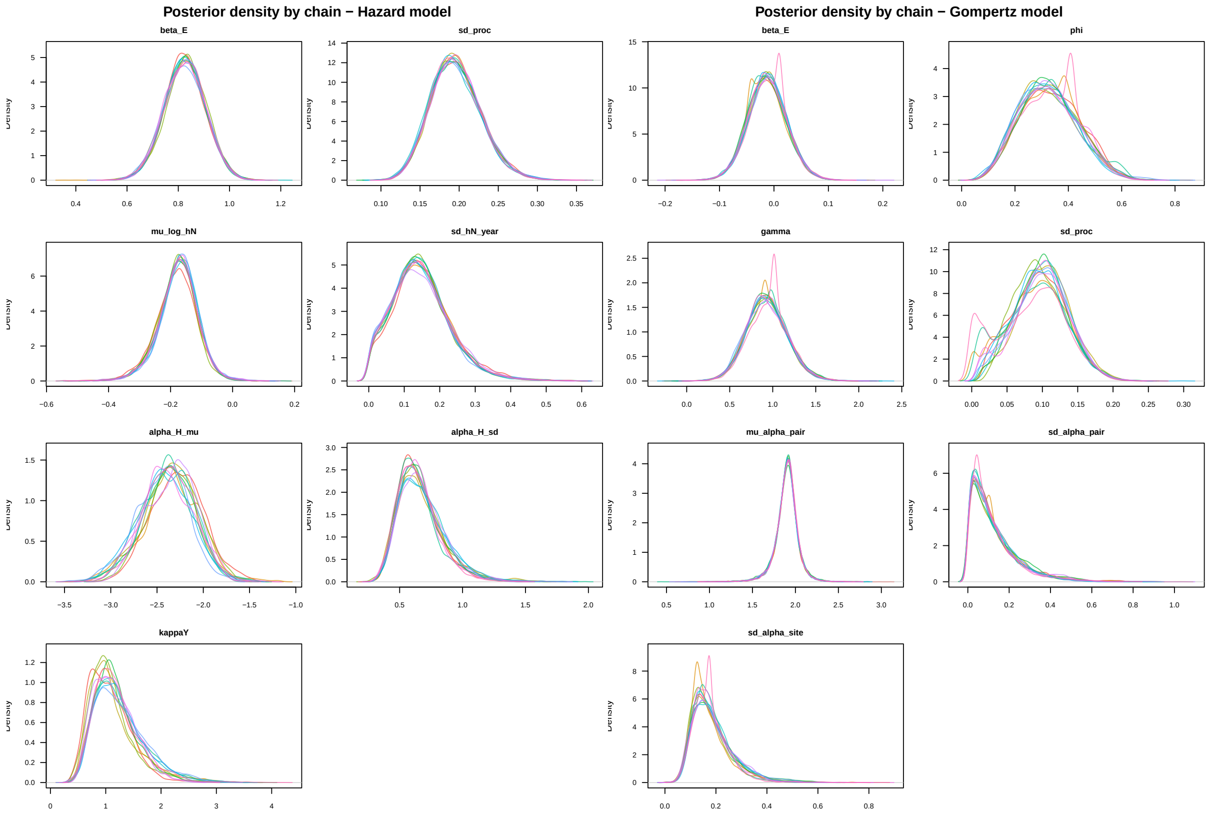


**(a)** Hazard Model

**(b)** Gompertz Model

**Figure 3:** Graphs showing density plots for all parameters in the Hazard and Gompertz models.

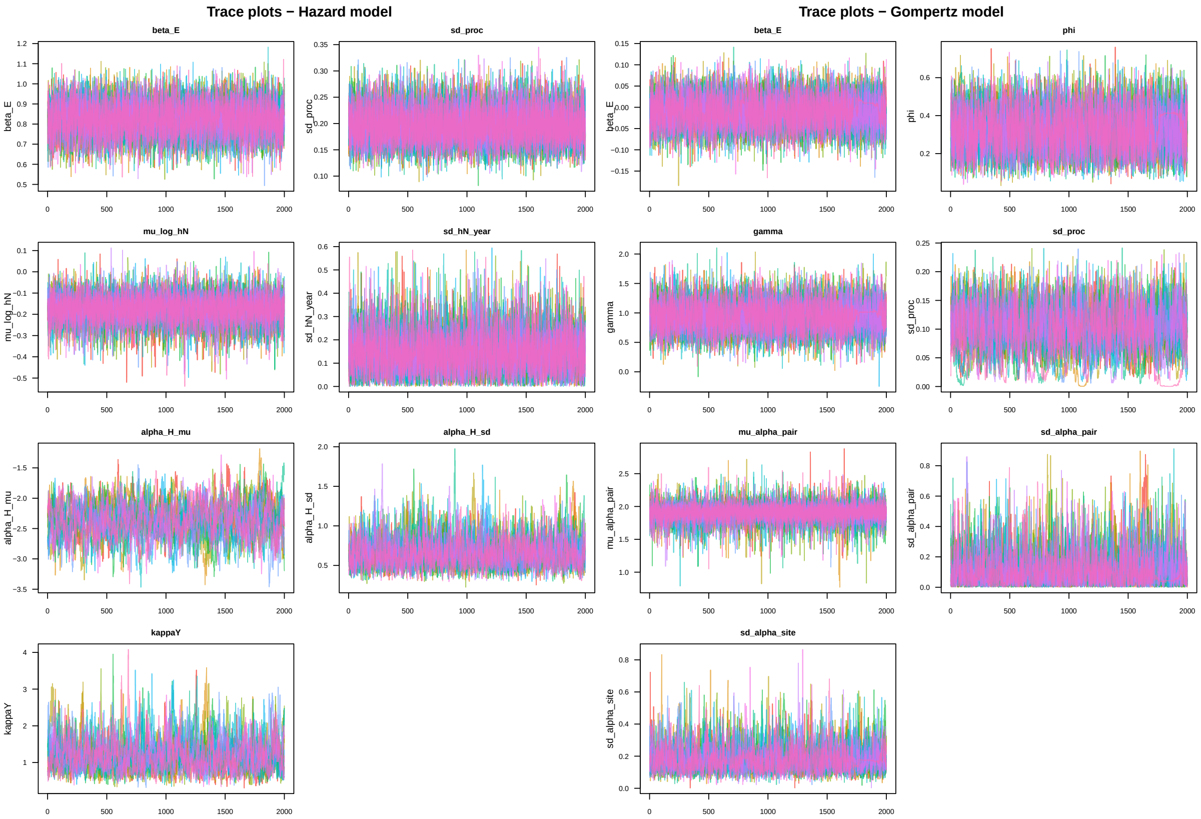


**(a)** Hazard Model

**(b)** Gompertz Model

**Figure 4:** Graphs showing trace plots for all parameters in the Hazard and Gompertz models.
